# A non-human primate model with Alzheimer’s disease-like pathology induced by hippocampal overexpression of human tau

**DOI:** 10.1101/2022.10.26.513832

**Authors:** Zhouquan Jiang, Jing Wang, Bin Luo, Fan Bai, Yongpeng Qin, Huiyi Wei, Shaojuan Zhang, Junjie Wei, Guoyu Ding, Long Ma, Shu He, Rongjie Chen, Lu Wang, Hao Xu, Xiangyu Wang, Gong Chen, Wenliang Lei

## Abstract

Alzheimer’s disease (AD) is one of the most burdening diseases of the century with no disease-modifying treatment yet. Non-human primates (NHPs) share genetic, anatomical and physiological similarities with humans, making them an ideal model for investigating the pathogenesis and therapeutics of AD. However, the applications of NHPs in AD research have been hindered by the paucity of spontaneous or induced monkey models for AD due to their long generation time, ethical considerations and technical challenges in making genetically modified monkeys. Here we developed an AD-like NHP model by overexpressing human tau in bilateral hippocampi of adult rhesus macaque monkeys. We evaluated the pathological features of these monkeys with immunostaining, cerebrospinal fluid (CSF) analysis, magnetic resonance imaging (MRI), positron emission tomography (PET) scan, and behavioral tests. We demonstrated that after hippocampal overexpression of human tau, the rhesus macaque monkeys displayed multiple pathological features of AD, including neurofibrillary tangle formation, neuronal loss, hippocampal atrophy, neuroinflammation, Aβ clearance deficit, blood vessel damage and cognitive decline. This work establishes a human tau-induced AD-like NHP model that may facilitate mechanistic studies and therapeutic treatments for AD.

## 1. Introduction

Alzheimer’s disease (AD) is the most prevalent neurodegenerative disorder worldwide with no curative treatment, which is partially due to the dismal track record of bench-to-bedside translation of intervention strategies developed and evaluated in AD rodent models (2021; Graham, et al. 2017). Non-human primates (NHPs) share close phylogenetic relationship as well as many physiological and pathological parallels with humans. They have been emerging as indispensable translational tools in the identification and validation of early diagnostic markers, as well as in the development and evaluation of safe and effective treatments for AD (Haque and Levey 2019; Vitek, et al. 2020). To the best of our knowledge, all the NHP models of AD available to date can be categorized into either spontaneous or artificially induced models, and the latter can be further classified as genetically or non-genetically modified models (Li, et al. 2019). Many NHPs including rhesus macaques, mouse lemurs and common marmosets exhibit Aβ deposits, incipient tau pathology and cognitive deficits as they grow old. Unfortunately, these naturally aged monkeys are not very consistent phenocopies of AD patients, since aging is essentially a different process than AD. Moreover, their limited availability, prohibitive breeding cost and very long generation period significantly restrict the broader application of spontaneous NHP models (Arnsten, et al. 2019; Geula, et al. 2002; Giannakopoulos, et al. 1997; Sani, et al. 2003). In accordance with a series of hypotheses of AD onset, NHP models induced by cholinergic neuron injury, injection, formaldehyde exposure, and streptozotocin injection each reflect certain pathological aspects or behavioral deficits of AD. However, there are continuing controversies regarding the validity of the theoretical foundations of these models. Meanwhile, uniformity of model development, side effects of drug administration, penetration damages caused by surgical procedures, and still substantial time frame considerably limit the potential utilization of non-genetically engineered NHP models (Forny-Germano, et al. 2014; Geula, et al. 1998; Melamed, et al. 2017; Park, et al. 2015; Zhai, et al. 2018). Today, a couple of genetic NHP models of AD, such as marmosets with edited *PSEN1* and *APP* transgenic cynomolgus monkeys, are looming on the horizon. Yet again, these promising models have their limitations. Ethical considerations, under-representation of sporadic AD, non-tissue specific gene expression, unpredictable genomic integration, and suboptimal gene transfer efficiency lead to the paucity and undermine the reliability of genetically modified NHP models (Sasaguri, et al. 2020; Seita, et al. 2020).

In order to remove the roadblocks preventing the wide applications of NHP models in AD research, we attempted to establish a convenient and reliable monkey model to facilitate preclinical studies on AD. Growingly recognized as a major driver of AD, tau captured our attention, since it surpasses Aβ in predicting the location of brain atrophy in AD patients, and correlates closely with AD progression and cognitive decline (Dujardin, et al. 2020; La Joie, et al. 2020; Morshed, et al. 2021; Qiu, et al. 2021). In fact, the rhesus monkeys who received injections of pathological tau-expressing adeno-associated virus (AAV) in unilateral entorhinal cortex display misfolded tau propagation accompanied by extensive microglial responses and robust alterations of cerebrospinal fluid and plasma biomarkers for AD (Beckman, et al. 2021). Therefore, in this study, we created a new Alzheimer’s-like NHP model by overexpressing human tau (hTau) in bilateral hippocampi of mid-aged rhesus monkeys. These AD-like monkeys can be generated through a single stereotaxic injection of AAVs within 2 months, yet they exhibit many defining pathological features of AD, such as neuronal loss, inflammatory responses, medial temporal lobe atrophy, elevated Aβ burden, neurofibrillary tangle formation, vascular abnormalities and multidomain cognitive dysfunctions. Once adopted by the research community, they are expected to add to the arsenal of AD NHP models and to contribute to the fight against the devastating Alzheimer’s disease.

## 2. Materials and methods

### 2.1. Reagents, AAVs and antibodies

The anesthetics, analgesics and sedatives we have used in this study includes: Zoletil™ 50 (Virbac, France), Dexdomitor^®^ (dexmedetomidine, Orion Pharma, Finland), propofol (Guangdong Jiabo Pharmaceutical, China), atropine (Shanghai Quanyu Biotechnology, China), lidocaine (Shandong Hualu Pharmaceutical, China). The primary antibodies we used in this study are as follows: chicken anti-GFP (Abcam, ab13970, 1:1000), rabbit anti-Tau (Dako, A0024, 1:1000), guinea pig anti-NeuN (Millipore, ABN90, 1:1000), mouse anti-pTau (Invitrogen, MN1020B, 1:500), rat anti-GFAP (Invitrogen, 13-0300, 1:1000), rabbit anti-Iba1 (Fujifilm, 019-19741, 1:1000), rabbit anti-CD68 (Abcam, ab125212, 1:500), mouse anti-Aβ (BioLegend, 800701, 1:500), rabbit anti-laminin (Sigma, L9393, 1:1000), rabbit anti-AQP4 (Proteintech, 16473-1-AP, 1:500), mouse anti-PECAM-1 (Thermo Fisher, MA5-13188, 1:1000). The secondary antibodies we used in this study are listed below: donkey anti-mouse, rat, rabbit, chicken, and guinea pig, Alexa Fluor 488, 555, and 647 (Invitrogen or Jackson ImmunoResearch, A11039, A21206, A21208, A31570, A31572, A21435, A21207, A31571, A21247, 712-165-150, 711-605-152, 706-605-148, 1:1000 or 1:500). The DAPI and Thioflavin-S were obtained from Roche (10236276001, 1:1000) and Sigma (T1892-25G, 1:5000), respectively. And all the AAVs we have used in this study were produced by PackGene Biotech Inc. (Guangzhou, China).

### 2.2. Animals

Nine adult rhesus macaques (Macaca mulatta, seven males and two females, 5 to 12 years old, body weights: 5~8 kg) were used in this study. The macaques were maintained in accordance with the standards set forth in the 8th edition of the *Guide for the Care and Use of Laboratory Animals* (NRC, 2011). All of them were housed in individual cages which can be easily sanitized and placed in climate-controlled rooms at Guangdong Yuan Xi Biotech Co., Ltd. (Guangzhou, China), which was fully accredited by the Association for Assessment and Accreditation of Laboratory Animal Care (AAALAC) International, under the close supervision of three laboratory animal technicians and the Institute veterinarians. All experimental procedures were approved by the Institutional Animal Care and Use Committee (IACUC) of Guangdong Yuan Xi Biotech Co., Ltd. and Jinan University and were in full compliance with the “Guide for the Care and Use of Laboratory Animals of the Institute of Laboratory Animal Science (est. 2006)” and “The use of non-human primates in research of the Institute of Laboratory Animal Science (est. 2006)” to ensure the safety of personnel and animal welfare.

### 2.3. Surgical procedures and stereotactic injection of AAVs

All the monkeys were fasted for ~12 hours prior to general anesthesia to help prevent vomiting and aspiration of stomach contents while under general anesthesia. atropine (0.02 mg/kg, intramuscular injection) was given preoperatively to reduces secretions in the mouth and respiratory passages of the monkeys. Anesthesia was induced by Zoletil™ 50 (tiletamine and zolazepam, 5–10 mg/kg, intramuscular) and Dexdomitor^®^ (dexmedetomidine, 10-20 mcg/kg, intramuscular). During surgery, anesthesia was maintained with propofol (2.5-3.5 mg/kg, intravenous), while lidocaine (1-4 mg/kg, subcutaneous, intramuscular) was also administered as a local anesthetic before the operation to further reduce pain.

For hippocampal AAV injection, the monkeys were placed on a heated V-top surgical table with their heads stabilized on a stereotaxic frame (RWD Life Science, Shenzhen, China). The precise positions (6 within each hippocampus) for stereotaxic injection were determined based on the T1-weighted MRI scans of the monkey brains. A hand-held cranial microdrill (RWD Life Science, Shenzhen, China) was used to drill several small holes in the skull to allow the injection needles to pass through. Then 7.5ul AAVs (10^11~12^ GC/ml, flow rate: 800 nl/min) were injected into each injection site using a 25 μl microsyringe (Hamilton Company, USA) controlled by a Micro4™ controller and an UltraMicroPump 3 (World Precision Instruments, USA). After each injection, the microsyringe needle was kept in place for 10 additional minutes before it was slowly withdrawn. During the surgical procedures, the blood oxygen level (> 95%), heart rate (150–220/min), respiratory rate (10–25/min) and blood pressure (> 60 mmHg mean value, and > 90 mmHg for systolic pressure) were all detected and collected with a pediatric vital signs monitor (JRTYL, Hunan, China). Finally, the muscles and skins around the wound were repeatedly cleaned and sutured, and after the operation, penicillin sodium (100,000 U/kg, intramuscular) was given for three consecutive days to prevent wound infection.

### 2.4. Immunohistochemistry

The monkeys were euthanized via sodium pentobarbital overdose (100 mg/kg), and transcardially perfused first with ice-cold PBS and later with a mixture of paraformaldehyde (4% PFA) and sucrose (10%). After perfusion, dissected out the whole monkey brains, sliced them into 1 cm coronal blocks with a monkey brain matrix (Shanghai Tow Intelligent Technology, China), and then post-fixed and dehydrated these blocks in 4% paraformaldehyde (48 hours) and gradient sucrose solutions (10, 20 and 30%). The dehydrated brain blocks were embedded in Tissue-Tek^®^ O.C.T. Compound (Sakura Finetek, USA), and then serially sectioned at the coronal plane on a cryostat (CryoStar™ NX50, Thermo Fisher Scientific, USA) at 50 μm thickness.

For immunofluorescence, free floating brain sections were first washed with PBS and blocked in blocking solution (5% normal donkey serum, 3% BSA and 0.5% Triton X-100 in PBS) for 1 hour at room temperature, and then incubated overnight at 4°C with primary antibodies diluted in blocking solution. After at least 3× washes with PBS, the brain sections were incubated with DAPI and appropriate secondary antibodies conjugated to Alexa Fluor 488, Alexa Fluor 555, or Alexa Fluor 647 for 2 hours at room temperature, followed by extensive washing with PBS. Finally, the brain sections were mounted with VECTASHIELD^®^ antifade mounting medium (VECTOR Laboratories, USA) and sealed with nail polish.

### 2.5. Wide-field and confocal fluorescence imaging

Most of the fluorescence images were acquired with a modular microscope user interface software (64 bit version of ZEN 2.5, Carl Zeiss, Jena, Germany) on either a conventional widefield epi-fluorescence microscope (Carl Zeiss Axio Imager Z2, Jena, Germany) or a confocal laser scanning microscope (Carl Zeiss LSM 880 with Airyscan, Jena, Germany).

### 2.6. Magnetic resonance imaging (MRI) and positron emission tomography (PET) scans

All MRI scans were performed on a 3.0T MRI scanner (Discovery MR750 3.0T, GE Healthcare, USA) equipped with an 8-channel customized head coil for macaques (Medcoil MK80, Suzhou, China) at the PET/CT-MRI center in the First Affiliated Hospital of Jinan University. Prior to each MRI scan, the monkeys were fasted for at least 6 hours and anesthetized with an intramuscular injection of Zoletil™ 50 and Dexdomitor^®^. When the MRI data were collected to determine the coordinates for stereotaxic injection of AAVs, a pair of glass capillary tubes (World Precision Instruments, USA) filled with glycerol were fixed onto the top of the skull ahead of the scans to serve as spatial reference points. The whole-brain images were acquired with a 3D Bravo T1 sequence (TR = 8.4 ms, TE = 3.5 ms, slice thickness = 0.5 mm, matrix size = 300 × 300, FOV = 15×15 cm), a CUBE T2 sequence (TR = 2500.0 ms, TE = 108.9 ms, slice thickness = 0.5, matrix size = 320 × 320, FOV = 15×15 cm), and a T2 FLAIR sequence (TR = 8400.0 ms, TE = 151.7 ms, slice thickness = 1.0 mm, matrix size = 256 × 256, FOV = 16×16 cm).

All ^18^F-FDG and ^18^F-T807 PET scans were conducted using a 128-slice time-of-flight PET/CT scanner (GE Discovery PET/CT 690 Elite, GE Healthcare, USA) at the PET/CT-MRI center in the First Affiliated Hospital of Jinan University. Before each PET scan, the macaques were fasted and anesthetized as previously mentioned in the last paragraph. The radiochemical purity of both ^18^F-FDG and ^18^F-T807 were over 99%. Each monkey was intravenously injected with ^18^F-FDG (approximately 150 MBq, 0.3~0.5 mCi/kg) or ^18^F-T807 (approximately 150 MBq, 0.3~0.5 mCi/kg) while awake via the posterior saphenous vein. After 50 min, the macaque was placed into the scanner. Head position was fixed with a stereotactic frame, followed by a static PET scan. Protocols: CT scan was performed firstly, followed by a ten-minute static positron emission data collection at 60 min post-injection. The PET data were attenuation-corrected by integrated CTAC technology. The PET/CT and MR images were co-registered and analyzed with software PMOD. The standard uptake value (SUV) of each region of interest (ROI), which were decay corrected back to the radioligand injection time point, were quantitatively extracted based on individual Atlas.

### 2.7. Animal behavioral studies

Delayed response (DR) task was conducted using the Wisconsin General Test Apparatus (WGTA), in which a monkey sat in its home cage in front of a tray that contains 3 food wells covered by identical swing-away lids. Initially, the experimenter baited one of the wells with food in front of the monkey, covered it, and then lowered an opaque screen to block the food tray from the monkey’s view. After a certain delay period the screen was removed, which allowed the monkey to retrieve the food from the baited food well which had to be recalled from working memory. Food reinforcers were randomly distributed among 3 wells over 30 trials per day (≥ 26 correct choices for 3 consecutive days to pass), and the length of the delays were gradually increased according to a stepwise procedure as the monkeys demonstrate mastery of the task.

Delayed matching-to-sample (DMTS) task was also performed with a WGTA. The experimenter first presented a sample visual stimulus (one of katakana characters or LEGO^®^ blocks) to the monkey. Following a randomly interval (up to 5 seconds), the same sample visual stimulus and another visual distractor were both presented to the monkey, and the monkey needed to choose the visual stimulus that matched the sample stimulus to get the food reward. Each incorrect choice was followed by a 10 seconds timeout. The monkeys took 30 DMTS trials per day (≥ 26 correct choices for 3 consecutive days to pass), and both the sample visual stimuli and the visual distractors were randomly selected each day from either 46 basic katakana alphabet letters or a large pool of LEGO^®^ blocks (> 200 combinations).

Rotating Brinkman board task is a manual dexterity task for monkeys, in which 14 food pellets were placed in 14 narrow slots (60□mm length ×□6□mm width□×□6□mm depth) on a rotating brinkman board (240□mm in diameter), and the monkeys were trained to retrieve all the food pellets from the slots using one of their hands. Monkeys were trained to master this task for 3 consecutive days before they were tested each day for retrieval accuracy and speed for 5 consecutive days. During both the training and testing sessions, the monkeys performed 4 sessions each day, among which the board was rotating clockwise in 2 sessions and counterclockwise in the other 2 sessions, and the monkeys were allowed to only use their left hands in 2 sessions and right hands in the other 2 sessions. All experimental procedures were recorded with FHD video cameras.

### 2.8. Single molecule array (Simoa)-based CSF analysis

To collect the CSF from the monkeys, a lumbar puncture was performed with a 22G traumatic needle between L4/L5, and the CSF samples were transferred to sterile low-binding microcentrifuge tubes, followed by centrifugation at 10000G at 4□°C for 5 min. Next, all the CSF samples were analyzed for the total-tau, phospho-tau 181, Aβ42 and Aβ40 peptides levels using Quanterix’ ultrasensitivity digital biomarker detection technology with the Simoa^®^ HD-X Analyzer™ (Quanterix, Billerica, MA, USA), the latest model fully automated Simoa bead-based immunoassay platform. The assay kits been used in this study were Simoa^®^ Neurology 3-Plex A Kit (101995-Lot 502593, Quanterix, USA) and Simoa™ pTau-181 Advantage V2 Kit (103714-Lot 502539, Quanterix, USA).

### 2.9. Statistical analysis

Image processing was performed using the 64 bit versions of ImageJ (National Institutes of Health, USA) and ZEN 2.5 (Carl Zeiss, Germany). The 3D reconstructions and tissue volumes were obtained using the Brainsight software (Rogue Research, Montréal, Québec, Canada).

All statistical analyses were performed with GraphPad Prism v8.4.2 (GraphPad Software, La Jolla, CA, USA), and datasets were assessed for normality parameters prior to the significance tests. Statistical significance was assessed using two-tailed Student’s t test when comparing two groups. When analyzing multiple groups, we used one-way ANOVA with Tukey’s post-hoc test to determine the statistical significance. For the variables measured longitudinally at several time points, we analyzed the data using repeated measures ANOVA with Tukey’s post-hoc test. A p value < 0.05 was considered significant.

Data are presented as mean ± SEM, and data were recorded and analyzed blindly, wherever possible.

## 3. Results

### 3.1. Establish a non-human primate model with Alzheimer’s-like pathology through hippocampal overexpression of human tau

To develop a tauopathy-induced non-human primate model with Alzheimer’s-like pathology, we employed adult rhesus monkeys (11-13 years old), and selected the hippocampus as the site of human tau overexpression since it is closely related to Alzheimer’s disease (Criscuolo, et al. 2017; Halliday 2017; Mu and Gage 2011). Under monitored anesthesia, we injected human tau-expressing AAVs in bilateral hippocampi of adult rhesus monkeys. After 4-11 weeks following viral injection, we evaluated these monkeys for changes in tau levels, neuronal population, neuroinflammation and blood vessels with immunostaining. Meanwhile, we also tested these monkeys for changes in tau levels, hippocampal mass, CSF AD biomarkers, cognitive and motor functions with MRI/PET imaging, CSF analysis and behavioral tests before and after the model construction (Fig. 1A).

**Figure 1.**
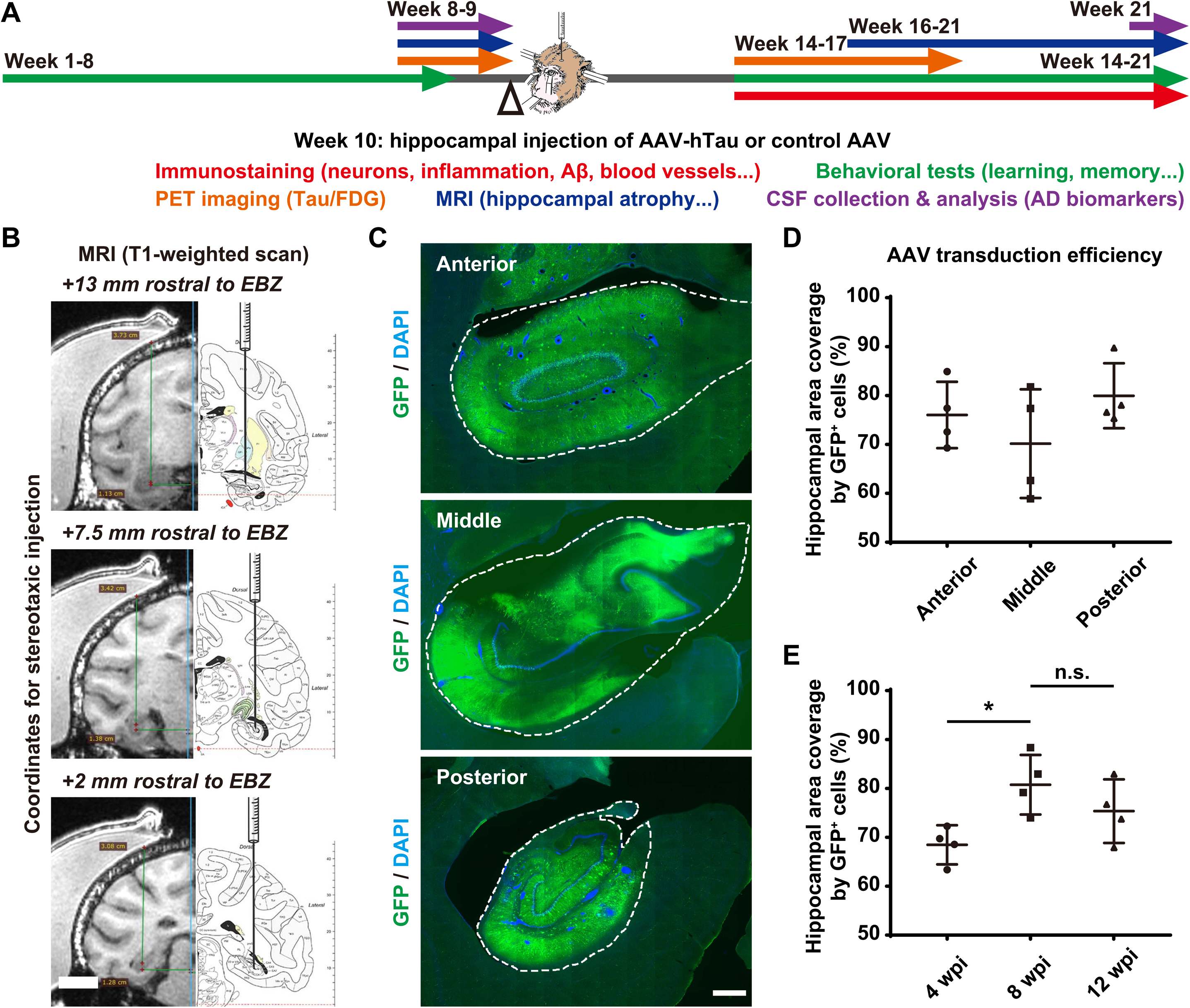
Overexpression of exogenous genes throughout bilateral hippocampi and construction of a non-human primate model with Alzheimer’s-like pathology by stereotactic injection of AAVs. **(A)** Schematic diagram depicts the construction and evaluation process of the rhesus monkey model with Alzheimer’s-like pathology. The stereotactic injection of AAVs is performed between week 9 and week 10. The monkeys are evaluated for changes in tau expression, neuronal survival, neuroinflammation, blood vessels, hippocampal volume, CSF biomarkers for AD, cognitive functions and fine motor skills before and after the viral infection. The whole process will take around 21 weeks. **(B)** The T1-weighted MRI scans and coronal atlas of the rhesus monkey brain illustrate the coordinates for stereotaxic injection of AAVs. Six sets of coordinates are determined for each individual animal before the viral injection based on 3-dimensional reconstruction of monkey brain from T1-weighted MRI scans. EBZ, ear-bar zero. **(C)** Representative images of GFP immunostaining show the AAV-induced broad expression of GFP (exogenous gene) throughout the monkey hippocampus 8 weeks after viral injection. The dashed lines outline the edges of the hippocampi. Scale bar, 1 mm. **(D-E)** Quantifications of the average transduction efficiencies of the AAVs in terms of hippocampal area coverage by GFP+ cells in different regions of the hippocampus (D) or at different time points after viral injection (E). *P < 0.05, and “n.s.” means “not (statistically) significant” (P>0.05). One-way ANOVA with Tukey’s post-hoc test. N = 4.

For each monkey, six sets of coordinates for stereotaxic injection of AAVs were predetermined based on the T1-weighted MRI scan of its brain collected using a customized head coil for macaques (Medcoil MK80, Suzhou, China, Fig. 1B). After a single round of AAV injection performed at 6 evenly-distributed injection sites, AAV vectors (Syn::GFP) were efficiently delivered to a major portion of the hippocampus, judging by the AAV-induced broad expression of GFP throughout the hippocampus (Fig. 1C). The mean transduction efficiency of the AAVs with regard to hippocampal area coverage by GFP^+^ cells was ~ 75% across different regions of the hippocampus (Fig. 1D) with a temporal profile which increased from 4 to 8 weeks, peaked at around 8 weeks, and plateaued at 12 weeks after viral injection (Fig. 1E), indicating a broad and long-lasting expression of exogenous genes throughout the monkey hippocampus.

### 3.2. Detection of high levels of total tau and phosphorylated tau throughout the viral infected NHP brain

Overexpression of human Tau was conducted through delivering a pair of AAV vectors (Syn::FLPo and CAG::FRT-hTau) into the monkey hippocampus. These AAVs will overexpress site-specific recombinase FLPo in neurons under the Synapsin promoter, and then induce robust hTau expression under the strong ubiquitous CAG promoter via FRT recombination (Fig. 2A). To assess human tau expression level in the viral infected hippocampus, we performed tau immunostaining on monkey brain slices collected 6 and 10 weeks after AAV injection. The expression of tau in the hippocampus of the monkeys infected with AAV Syn::FLPo and AAV CAG::FRT-hTau is significantly higher compared to the control monkeys (Fig. 2B, top row), and excessive tau accumulation is evident in both somas and apical dendrites of many hippocampal neurons (Fig. 2B and Supplementary Fig. 1). Quantitatively, hippocampal tau levels at 6 and 10 weeks after viral injection were 3-4 folds of those in the control monkeys (Fig. 2C).

**Figure 2.**
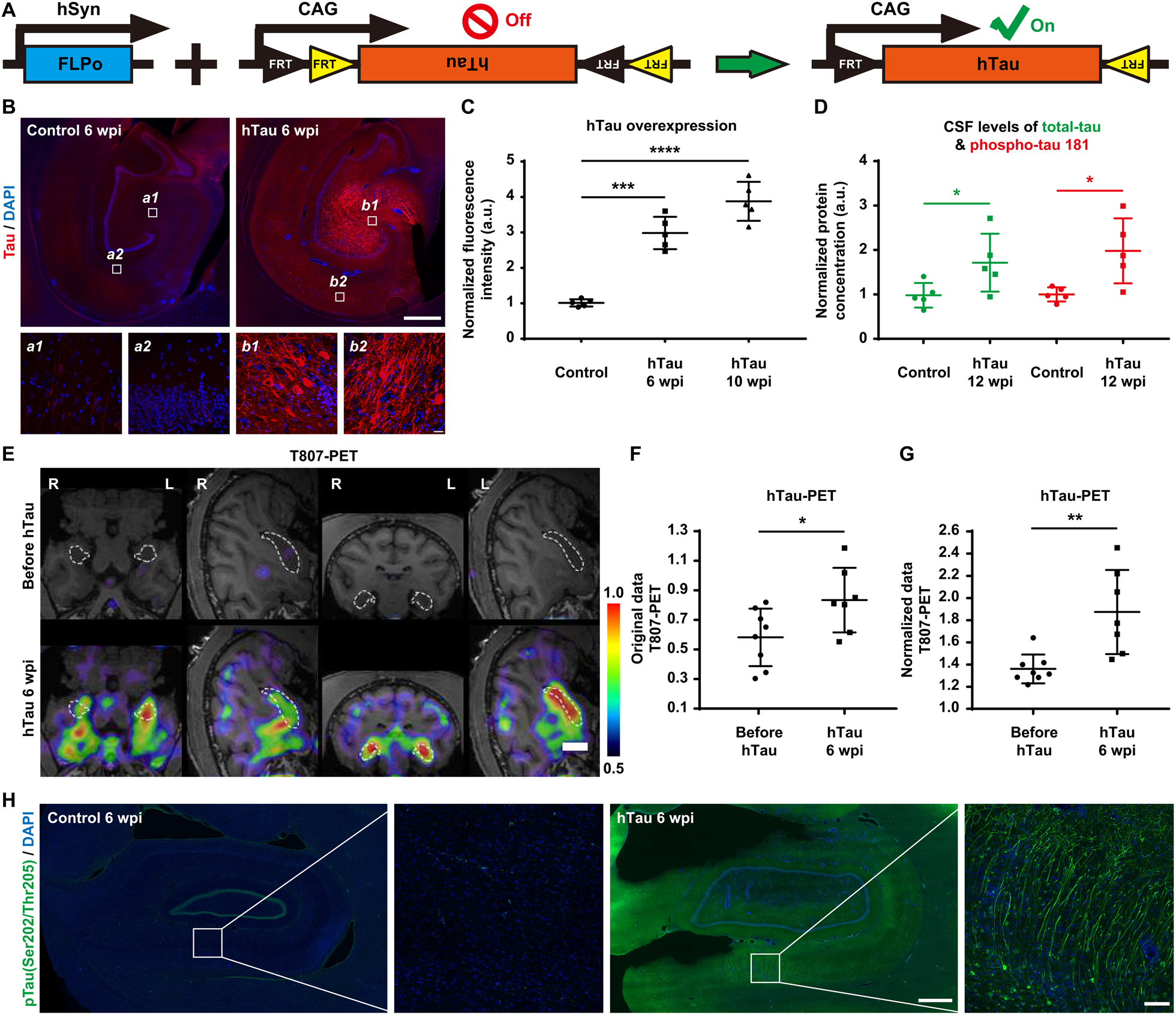
High levels of total tau and phosphorylated tau throughout the monkey hippocampus after AAVs injection. **(A)** Schematic diagram of our engineered AAV constructs (Syn::FLPo and CAG::FRT-hTau) used to express FLPo recombinase, which in turn will activate the expression of hTau. **(B)** Representative images of tau immunostaining demonstrate the AAV-induced overexpression of tau in neurons within the monkey hippocampus 6 weeks after viral injection. Compared to that of endogenous tau (left), the expression level of tau in the hTau overexpressed monkey brains (right) is significantly higher, especially in neuronal somas and apical dendrites. Insets show higher magnification of the CA3 (a1, b1) and CA1 (a2, b2) regions. Scale bar, 1 mm (top) and 20 μm (inset, bottom). **(C)** Quantification of the normalized fluorescence intensity of tau staining in the monkey brains. Expression levels of tau in the monkey hippocampus 6 and 10 weeks after hTau overexpression are significantly higher than that of endogenous tau in the control monkey hippocampus. ***P < 0.001; ****P < 0.0001. One-way ANOVA with Tukey’s post-hoc test. N = 5. **(D)** Quantification of the normalized total-tau (green) and phospho-tau 181 (red) concentrations in monkey CSF determined by the SIMOA-based biomarker analysis. The CSF levels of total-tau and phospho-tau 181 are significantly higher in the hTau overexpressed monkey 12 weeks after AAVs injection. *P < 0.05. Student’s t-test. N = 5. **(E)** Representative ^18^F-T807 PET/MRI fusion images indicate higher tau expression in monkey hippocampus 6 weeks after AAVs injection. The linear color scale with the standardized uptake value (SUV) range is shown in the right part of the figure. The dashed lines outline the edges of the hippocampi. Scale bar, 1 cm. **(F-G)** Quantifications of hippocampal ^18^F-T807 retention (F, original SUV values; G, hippocampus-to-cerebellum SUV ratios) 6 weeks after hTau overexpression. *P < 0.05; **P < 0.01. Student’s t-test. N = 7. **(H)** Representative images of phospho-tau (Ser202, Thr205) immunostaining demonstrate the dramatic increase of phospho-tau in the monkey hippocampus 6 weeks after AAV-induced tau overexpression. The result suggests the accumulation of paired helical filaments (PHFs), which are the building blocks of the neurofibrillary tangles in Alzheimer’s brain. Insets show higher magnification of the CA1 regions. Scale bar, 1 mm (left) and 20 μm (right, inset).

Along with total-tau, phospho-tau 181 in plasma or blood correlates positively with disease severity of Alzheimer’s disease, and hence can serve as a specific biomarker for Alzheimer’s disease (Andreasen, et al. 2003; Karikari, et al. 2020; Mielke, et al. 2018). Therefore, we applied the single molecule array (Simoa) technique to detect the concentrations of total-tau and phospho-tau 181 in monkey CSF samples gathered before and after 12 weeks of hTau overexpression (Fig. 2D). As expected, both the amounts of total-tau and phospho-tau 181 in monkey CSF samples were markedly elevated 12 weeks after AAV injection, mimicking the characteristics of Alzheimer’s disease.

Recently, the radiotracer ^18^F-T807 has been widely used for PET scanning of the brain to help identify the presence and estimate the distribution of tau pathology, a distinctive characteristic of AD. Distinct from the traditional immunostaining approach, ^18^F-T807 PET imaging can be employed to repeatedly measure tau levels in the brain of the same animal over the course of weeks or even months (Passamonti, et al. 2017; Sander, et al. 2016; Scholl, et al. 2017). Consistent with the immunostaining and CSF analysis results, the ^18^F-T807 PET scan also exhibited significant tau accumulation in the monkey hippocampus 6 weeks after AAV injection (Fig. 2E). Both the original values of hippocampal ^18^F-T807 retention and the hippocampus-to-cerebellum ^18^F-T807 retention ratios increased considerably 6 weeks after hTau overexpression (Fig. 2F-G).

The hyperphosphorylation of tau leads to the formation of neurofibrillary tangles (NFTs) in Alzheimer’s disease. More specifically, the phosphorylation at S202 and T205 located in a proline-rich region of the tau molecule was reported to both enhance tau polymerization and induce tau filament formation (Moloney, et al. 2021; Neddens, et al. 2018). Here we tried to label pre-and mature NFTs by immunohistochemistry using phospho-tau (S202/T205) monoclonal antibody (AT8). Unexpectedly, after only 6 weeks of AAV-mediated hTau overexpression, many hippocampal neurons already exhibited strong positive staining for phospho-tau (S202/T205) in both somata and neuronal processes, indicating very rapid induction of NFT formation in the hippocampi of these monkeys (Fig. 2H).

More interestingly, the overexpression of tau could be detected throughout the monkey cortex 50 weeks after viral injection (Supplementary Fig. 2). This brain-wide spreading of tau pathology is possibly achieved through tau propagation using prion-like mechanisms (Beckman, et al. 2021; Colin, et al. 2020). Together, these results indicate that our hTau overexpression strategy has successfully generated hyperphosphorylated tau as well as tau tangles in the monkey hippocampus and cortex.

### 3.3. Significant neural degeneration induced by human tau overexpression

Progressive hippocampal neurodegeneration is one of the most prominent pathological features of Alzheimer’s disease, and strongly correlate with cognitive impairment in many Alzheimer’s patients (DeTure and Dickson 2019; West, et al. 1994). Therefore, we performed NeuN immunostaining to label the nuclei of post-mitotic neurons in the brain slices obtained from rhesus monkeys after hTau overexpression. Compared to the control group without hTau overexpression, we observed a substantial loss of NeuN immunoreactivity across the monkey hippocampus after 6 and 10 weeks of hTau overexpression (Fig. 3A, Supplementary Fig. 3).

**Figure 3.**
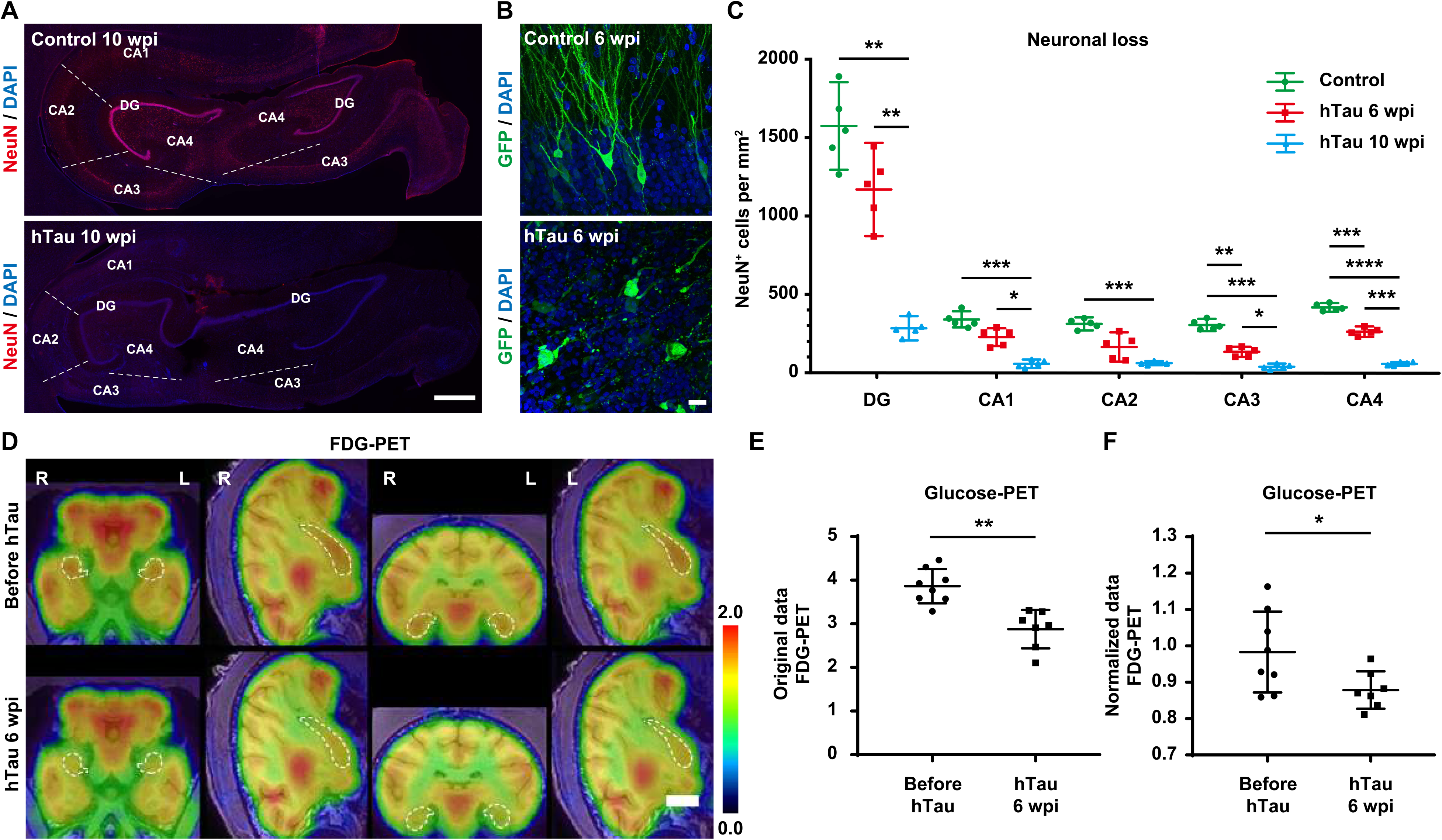
Dramatic neuronal degeneration and loss in the monkey hippocampus after hTau overexpression. **(A)** Representative images of NeuN immunostaining indicate neuronal degeneration and loss within the monkey hippocampus. The number of NeuN^+^ cells decreases significantly in the monkey hippocampus after 10 weeks of hTau overexpression. The dashed lines separate neighboring regions of the hippocampus. Scale bar, 1 mm. **(B)** Representative images of GFP immunostaining show marked neuritic dystrophy and neuronal damage in the monkey hippocampus after 6 weeks of hTau overexpression. Scale bar, 20 μm. **(C)** Quantification of the number of NeuN^+^ cells per mm^2^ in each hippocampal region of the monkeys without (green) or with 6 weeks (red) or 10 weeks of hTau overexpression (blue). *P < 0.05; **P < 0.01; ***P < 0.001; ****P < 0.0001. One-way ANOVA with Tukey’s post-hoc test. N = 5. **(D)** Representative ^18^F-FDG PET/MRI fusion images show a decreased hippocampal glucose metabolism, suggesting pronounced neuronal degeneration or death in monkey hippocampus 6 weeks after AAVs injection. The linear color scale with the standardized uptake value (SUV) range is shown in the right part of the figure. The dashed lines outline the edges of the hippocampi. Scale bar, 1 cm. **(E-F)** Quantifications of hippocampal ^18^F-FDG retention (E, original SUV values; F, hippocampus-to-cerebellum SUV ratios) 6 weeks after hTau overexpression. *P < 0.05; **P < 0.01. Student’s t-test. N = 7.

Considering that reduced NeuN immunoreactivity can be caused by not only neuronal loss but also depletion of the protein or loss of its antigenicity (Ünal-Çevik, et al. 2004), we analyzed the morphology of the neurons in monkey hippocampi with or without hTau overexpression. Six weeks after viral injection, the neurons in cornu ammonis regions of the monkey hippocampi infected with AAV CAG::FRT-hTau and AAV Syn::GFP exhibited the typical morphology of pyramidal neurons, whereas the same group of neurons infected with AAV Syn::FLPo, AAV CAG::FRT-hTau and AAV Syn::GFP showed degenerative features with disintegrated cellular structures (Fig. 3B), suggesting that the weak NeuN labeling is indeed closely correlated with neuronal degeneration and death. Quantitatively, the densities of NeuN^+^ cells in each hippocampal region (DG/CA1/CA2/CA3/CA4) of the monkeys declined by roughly 50~75% at 6 weeks and 80~90% at 10 weeks after hTau overexpression (Fig. 3C).

FDG-PET brain imaging is a very sensitive and powerful tool to detect subtle glucose metabolic changes in the brains, and hypometabolism in certain brain regions highly correlates with the pathological diagnosis of AD (Chetelat, et al. 2003; Herholz, et al. 2002; Mosconi, et al. 2009). Consistent with the NeuN^+^ immunostaining result, the metabolic rate for glucose in the hippocampal area of the monkey brains decreased significantly at 6 weeks after hTau overexpression (Fig. 3D). Both the original values of hippocampal ^18^F-FDG retention and the hippocampus-to-cerebellum ^18^F-FDG retention ratios dropped substantially 6 weeks after hTau overexpression, indicating marked neuronal degeneration throughout the monkey hippocampus (Fig. 3E-F).

### 3.4. Hippocampal atrophy following human tau overexpression

Reduced hippocampal volume can lead to profound amnesia, which is a core feature of Alzheimer’s disease. Furthermore, bilateral hippocampal atrophy is associated with increased risk for the progression of mild cognitive impairment (MCI) to Alzheimer’s disease (Apostolova, et al. 2006; Dubois, et al. 2014; Halliday 2017). Therefore, we tried to find out whether the monkey hippocampal volume changed after hTau overexpression. We utilized T1-weighted MRI brain scan to monitor the volume change of the hippocampus after hTau overexpression, and detected significant shrinkage of the hippocampus in the same monkey brain 12 weeks after viral injection (Fig. 4A).

**Figure 4.**
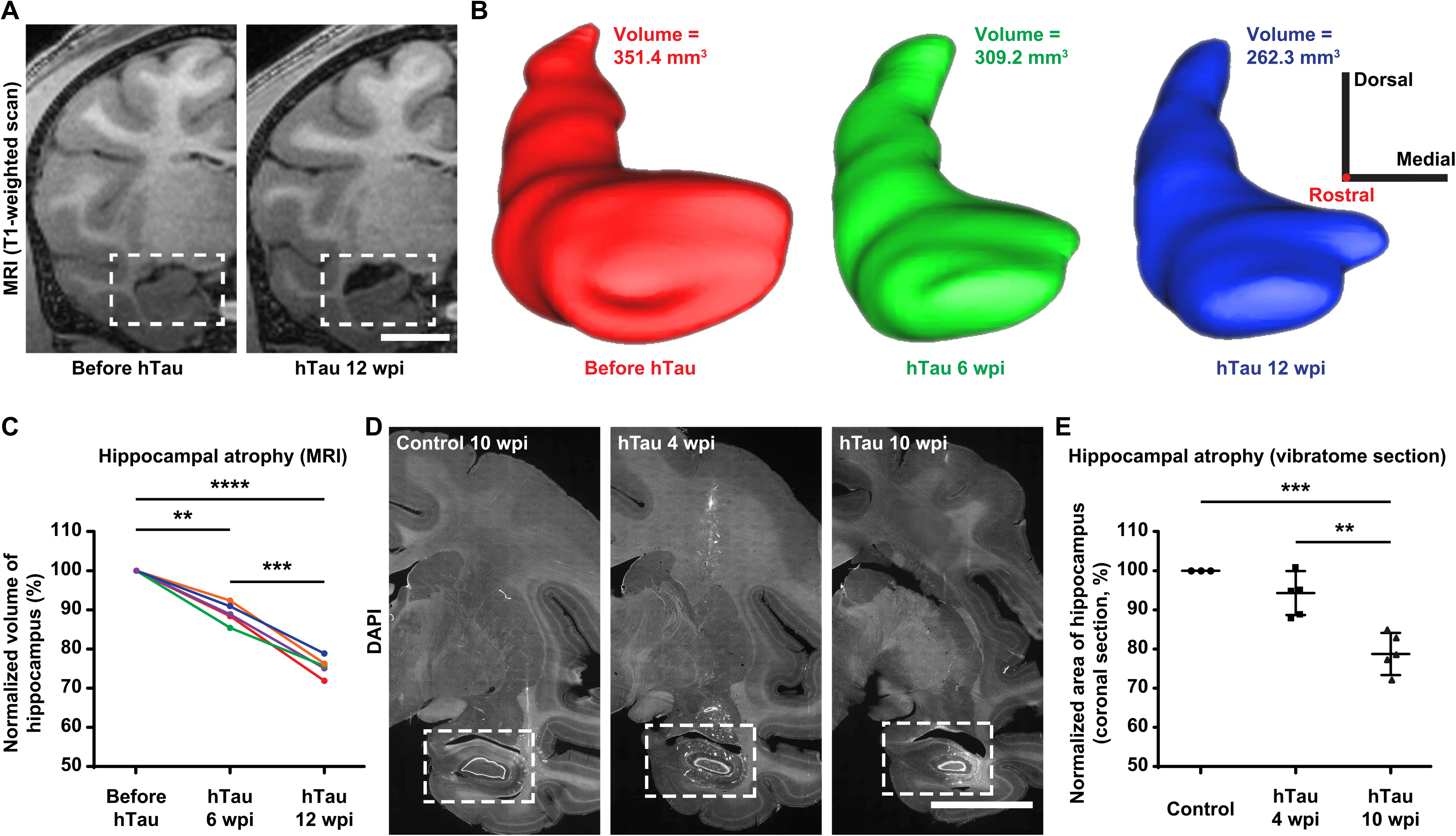
Substantial and progressive hippocampal atrophy after hTau over-expression. **(A)** Representative coronal T1-weighted MRI images demonstrate the hippocampal atrophy in monkey brains after 12 weeks of hTau overexpression. The hippocampal area measured in the coronal section shrinks significantly in the monkey brain 12 weeks after viral injection (right). The dashed boxes highlight the hippocampal areas. Scale bar, 1 cm. **(B)** Three-dimensional surface reconstructions and the volumes of the right hippocampus of a monkey at 3 different time points throughout the course of hTau overexpression (red, before AAVs injection; green, 6 weeks post injection; blue, 12 weeks post injection). This longitudinal study clearly reveals the progressive hippocampal volume decrease after hTau overexpression. The 3D reconstructions and tissue volumes are obtained using the Brainsight software (Rogue Research). 3D scale bar, 1 cm in each direction; the rostral-caudal axis is perpendicular to the screen with the rostral direction points out of the screen. **(C)** Quantifications of the longitudinal changes in the normalized hippocampal volumes defined by MRI scans. Each color represents one animal. **P < 0.01; ***P < 0.001; ****P < 0.0001. Repeated measures ANOVA with Tukey’s post-hoc test. N = 5. **(D)** Representative images of coronal vibratome sections (DAPI labeling) show the hippocampal atrophy in monkey brains after 4 and 10 weeks of hTau overexpression. The hippocampal area measured in the coronal section shrinks significantly in the monkey brain 4 (middle) and 10 (right) weeks after viral injection. The dashed boxes highlight the hippocampal areas. Scale bar, 1 cm. **(E)** Quantifications of the longitudinal changes in the normalized hippocampal areas measured in coronal vibratome sections. **P < 0.01; ***P < 0.001. One-way ANOVA with Tukey’s post-hoc test. N = 5.

By employing the Brainsight neuronavigation system, we reconstructed MRI sections of the monkey hippocampi into 3-dimensional models and measured the volumes at different time points after hTau overexpression. The results showed a progressive decrease in the hippocampal volume at 6 and 12 weeks after AAV injection, suggesting tauopathy-induced hippocampal atrophy (Fig. 4B). The quantification of normalized hippocampal volumes measured from the MRI scans exhibited roughly 10% decline at 6 weeks and 25% decline at 12 weeks in the size of hippocampal mass after hTau overexpression (Fig. 4C).

To corroborate with the MRI imaging analysis, we further analyzed the hippocampal volume by comparing the cross-sectional areas of the hippocampi in coronal vibratome sections of the monkey brains before and after hTau overexpression. Such a comparison revealed considerable and simultaneous hippocampal shrinkage and lateral ventricle enlargement both 4 and 10 weeks after viral injection (Fig. 4D). In terms of the extent of this hippocampal atrophy, the normalized hippocampal areas calculated from the coronal vibratome sections decreased by around 7% after 4 weeks and 20% after 10 weeks of hTau overexpression (Fig. 4E). Therefore, hTau overexpression results in a significant hippocampal atrophy.

### 3.5. Neuroinflammation induced by human tau overexpression

In Alzheimer’s disease astrocytes become reactive (activated), and reactive astrocytes are often found in close association with the Aβ plaques in the brains of AD patients. The concomitant reactive gliosis is involved in excessive neuroinflammation, oxidative stress, and neuronal dysfunction in AD. Such astrocyte activation can be either harmful due to loss of the neurotrophic effects and overproduction of proinflammatory molecules, or helpful because of attenuation of Aβ plaque growth and reduction in dystrophic neurites (Gonzalez-Reyes, et al. 2017; Kraft, et al. 2013; Pekny and Nilsson 2005; Perez-Nievas and Serrano-Pozo 2018).

We conducted immunostaining of GFAP, a hallmark of astrocytes, and discovered a substantial increase in GFAP signal across the monkey hippocampus after hTau overexpression (Fig. 5A, Supplementary Fig. 4). Many GFAP^+^ cells in the hippocampus underwent typical morphological changes seen in reactive astrocytes, such as cytoskeletal hypertrophy and outgrowth of particularly long processes (Fig. 5B). Quantitatively, hTau overexpression increased the density of GFAP^+^ cells by 2 to 4-fold in each region of the monkey hippocampus (Fig. 5C).

**Figure 5.**
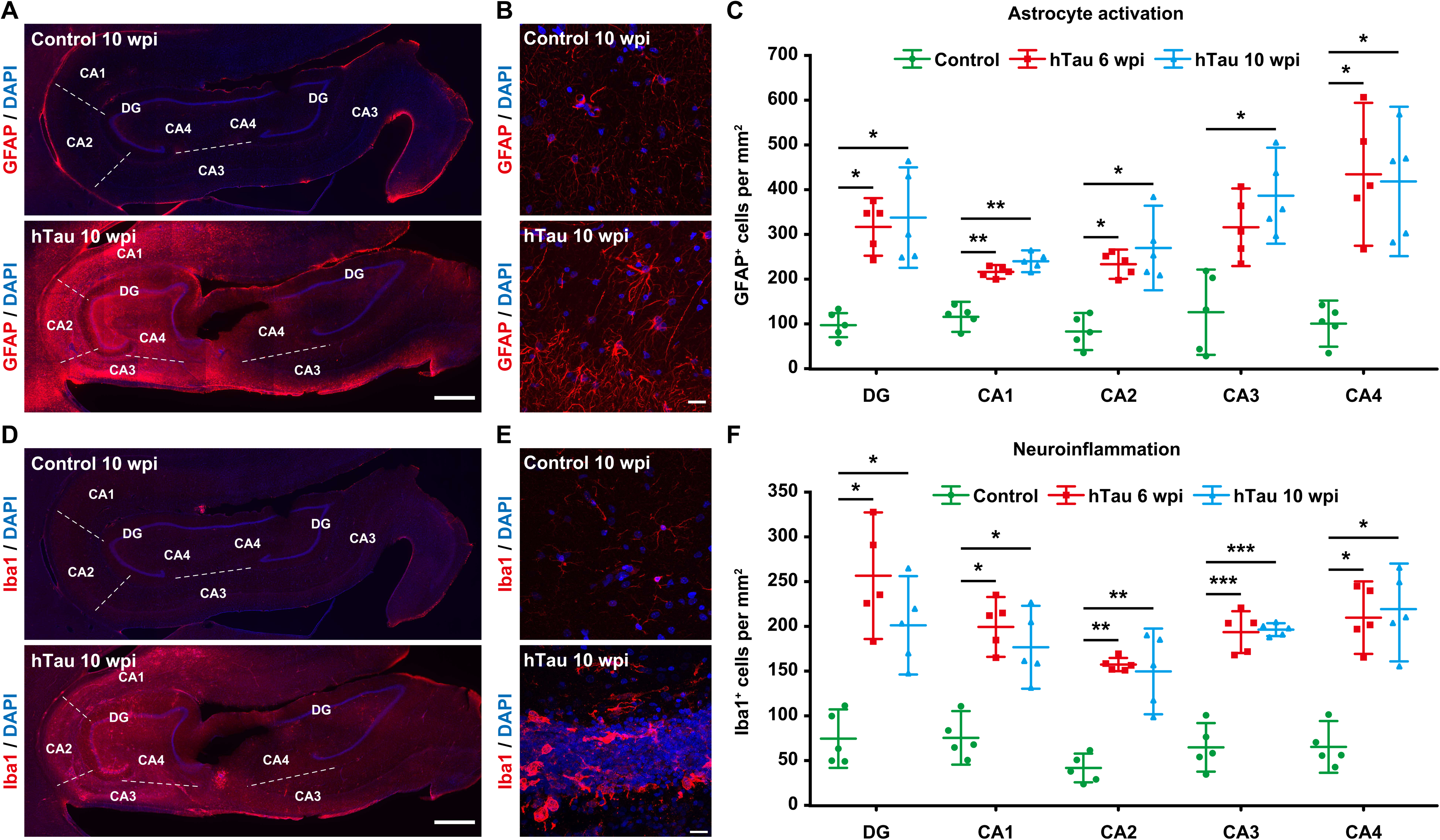
Extensive astrocyte activation and neuroinflammation in the monkey hippocampus after hTau overexpression. **(A)** Representative images of GFAP immunostaining demonstrate astrocytic activation within the monkey hippocampus. The number of GFAP^+^ cells increases significantly in the monkey hippocampus after 10 weeks of hTau overexpression. The dashed lines separate neighboring regions of the hippocampus. Scale bar, 1 mm. **(B)** Representative images of GFAP immunostaining show the transition of astrocytes from a resting state to a reactive state in the monkey hippocampus after 10 weeks of hTau overexpression. Scale bar, 20 μm. **(C)** Quantification of the number of GFAP^+^ cells per mm^2^ in each hippocampal region of the monkeys without (green) or with 6 weeks (red) or 10 weeks of hTau overexpression (blue). *P < 0.05; **P < 0.01. One-way ANOVA with Tukey’s post-hoc test. N = 5. **(D)** Representative images of Iba1 immunostaining suggest microglial activation and local neuroinflammatory response within the monkey hippocampus. The number of Iba1^+^ cells increases significantly in the monkey hippocampus after 10 weeks of hTau overexpression. The dashed lines separate neighboring regions of the hippocampus. Scale bar, 1 mm. **(E)** Representative images of Iba1 immunostaining show the transition of microglia from a highly ramified resting state to a less ramified, amoeboid reactive state in the monkey hippocampus after 10 weeks of hTau overexpression. Scale bar, 20 μm. **(F)** Quantification of the number of Iba1^+^ cells per mm^2^ in each hippocampal region of the monkeys without (green) or with 6 weeks (red) or 10 weeks of hTau overexpression (blue). *P < 0.05; **P < 0.01; ***P < 0.001. One-way ANOVA with Tukey’s post-hoc test. N = 5.

Microglia activation and neuroinflammation are considered as prominent factors contributing to the pathogenesis of AD. Genome-wide association studies have also demonstrated that many AD risk genes are highly expressed in microglia. Activated microglia can protect against AD by phagocytosing Aβ and cellular debris, or alternatively, aggravate AD by mediating synapse loss, exacerbating tau phosphorylation, impacting CNS homeostasis and secreting inflammatory factors (Hansen, et al. 2018; Heneka, et al. 2015; Leng and Edison 2021; Sarlus and Heneka 2017).

We performed immunostaining against Iba1, whose expression is up-regulated in activated microglia, and observed a marked increase of Iba1 immunoreactivity across the monkey hippocampus after hTau overexpression (Fig. 5D and Supplementary Fig. 5). In addition, many Iba1^+^ cells in the hippocampus transformed from a ramified, star-shape morphology into small, spherical, rod-shape or amoeboid-like morphologies (Fig. 5E). Moreover, the density of Iba1^+^ cells increased by 2 to 4-fold in each hippocampal region of the monkeys after hTau overexpression (Fig. 5F). Given that Iba1 is also expressed by resting microglia, whereas CD68 is commonly considered a marker of activated phagocytic microglia since it labels their lysosomes (Walker and Lue 2015), we also carried out immunostaining against CD68 in the hippocampi of the hTau-expressing monkeys. Elevated CD68 signals across the hippocampus further confirmed robust immune activation and neuroinflammation throughout the hippocampus of the NHP model of AD (Supplementary Fig. 6). Taken together, hTau overexpression can induce astrocytic and microglial pathologies in monkey hippocampus.

### 3.6. Decreased clearance and increased accumulation of Aβ after overexpressing human tau

Along with tauopathy, the production and deposition of Aβ is also a key feature of Alzheimer’s disease, and diverse lines of evidence suggests synergistic effects between Aβ and tau pathologies (Busche and Hyman 2020; Murphy and LeVine 2010). Furthermore, most patients with MCI or AD have reduced concentrations of Aβ peptide in the CSF, and lower CSF or plasma Aβ42/Aβ40 ratio has been considered as an indicator for greater risk of MCI or AD (Andreasen, et al. 1999; Graff-Radford, et al. 2007). After 10 weeks of hTau overexpression, we conducted immunostaining against Aβ, and detected some intracellular Aβ accumulation and extracellular Aβ plaque-like structures within many hippocampal slices of the monkey brains (Fig. 6A). Besides that, Thioflavin-S staining also revealed scattered Aβ plaque-like deposits surrounded by reactive microglia and reactive astroglia in many hippocampal slices of the monkey brains (Fig. 6B). Meanwhile, it is worth mentioning that the total quantity of the Aβ load was relatively small, possibly due to the relatively long period of time needed to develop Aβ pathologies throughout the brain. More interestingly, Simoa-based biomarker analysis showed that the CSF Aβ42 peptide level and Aβ42/Aβ40 ratio dropped considerably 12 weeks after hTau overexpression (Fig. 6C-D), suggesting a diminished capacity for Aβ clearance in these Alzheimer’s like monkeys.

**Figure 6.**
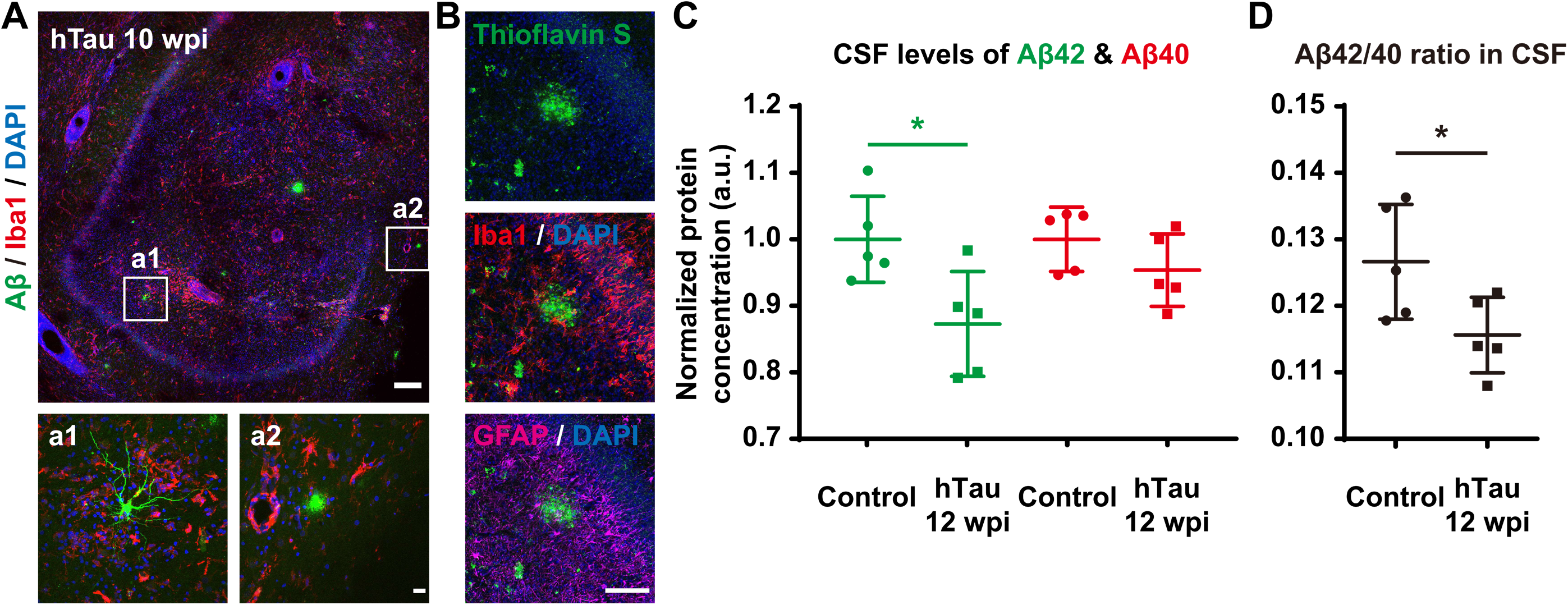
Decreased clearance and increased accumulation of Aβ throughout the monkey hippocampus after hTau overexpression. **(A)** Representative images of Aβ and Iba1 immunostaining suggest the development of Aβ pathology including intracellular Aβ oligomers and extracellular Aβ plaques within the monkey hippocampus 10 weeks after viral injection. Higher magnification insets show intracellular accumulation of soluble Aβ (a1) and extra-cellular plaque-like structure of Aβ aggregation (a2). Note the Iba1^+^ microglia surrounding the Aβ accumulating cells and plaques. Scale bar, 200 μm (top) and 20 μm (inset, bottom). **(B)** Representative images of Thioflavin-S staining labeling fibrillar Aβ deposits indicate the formation of Alzheimer’s Aβ plaques in the monkey hippocampus 10 weeks after AAV injection. Note the Iba1^+^ microglia and GFAP^+^ reactive astrocytes gathering around the Aβ plaque. Scale bar, 100 μm. **(C-D)** Quantification of Aβ42 (C, green) and Aβ40 (C, red) peptide concentrations, as well as Aβ42/Aβ40 ratio (D) in monkey CSF determined by the SIMOA-based biomarker analysis. The Aβ42 level and Aβ42/Aβ40 ratio in CSF are significantly higher in the hTau overexpressed monkey 12 weeks after AAVs injection. *P < 0.05. Student’s t-test. N = 5.

### 3.7. Vascular abnormality after human tau overexpression

Considering the clinical and pathological overlap between cerebrovascular disease and Alzheimer’s disease, many recent studies have shown that vascular dysfunctions including chronic brain hypoperfusion, blood-brain barrier leakage, and chronic vascular inflammation serve as a major trigger in the progression of sporadic AD. In fact, clinicopathological data also indicate an alarming positive feedback loop among vascular, glial and neuronal dysfunctions dictating the onset and development of the Alzheimer’s disease through its entire course (Attems and Jellinger 2014; de la Torre 2002; de la Torre 2004; Govindpani, et al. 2019). Immunostaining against laminin, a main component of the vascular basement membrane (Figure 7A), in monkey brain slices demonstrated vascular basement membrane thickening after hTau overexpression, which is one of the distinct changes of the blood vessels in AD (Thomsen, et al. 2017). Immunostaining against laminin also revealed many vascular abnormalities like string vessels (1~2 μm wide connective tissue strands, remnants of capillaries, that do not carry blood flow, Figure 7B, open arrows), tortuosity and bulging blood vessels (Figure 7B, closed arrowheads), and vascular occlusion (Figure 7B, open arrowheads) after hTau overexpression. Moreover, immunostaining against PECAM-1, a molecule expressed on endothelial and other vascular compartment cells, showed significant vascular damages and degeneration (Figure 7C) characterized by blood vessel ruptures (Figure 7D, arrows) and disintegration (Figure 7D, arrowheads) after hTau overexpression. Furthermore, immunostaining against AQP4, a water channel expressed at astrocyte endfeet, exhibited alteration in polarized distribution of AQP4 (Figure 7E, closed arrowheads & arrows) and astrocytic endfeet retraction from the blood vessels (Figure 7E, open arrowheads & arrows) after hTau overexpression. In terms of the extent of the vascular abnormalities or damages, around 25% and 45% of the capillaries displayed basement membrane thickening or some abnormal morphological alterations (string vessels / tortuosity and bulging vessels / vascular occlusion) after 6 and 10 weeks of hTau overexpression (Figure 7F). Meanwhile, around 25% and 35% of the capillaries were damaged or degenerated after 6 and 10 weeks of hTau overexpression (Figure 7G). Together, these results show that hTau overexpression can induce many vascular abnormalities and damages associated with AD in monkey brains.

**Figure 7.**
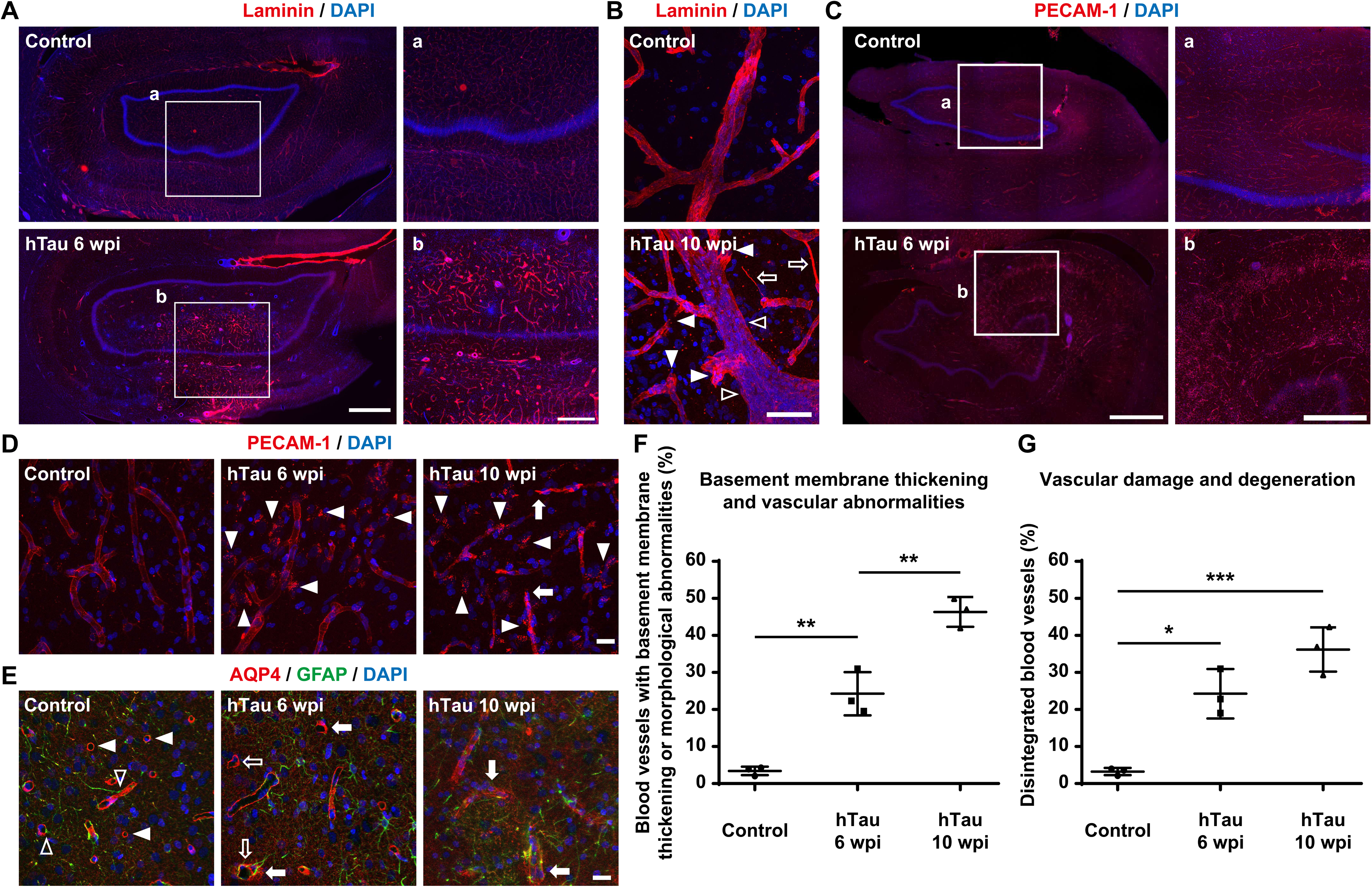
A variety of vascular abnormalities and damages throughout the monkey hippocampus after hTau overexpression. **(A)** Representative images of laminin immunostaining indicate vascular basement membrane thickening in the monkey hippocampus after 6 weeks of hTau overexpression. Insets show higher magnification images of the blood vessels in the hippocampus. Scale bar, 1 mm (left) and 500 μm (inset, right). **(B)** Representative images of laminin immunostaining indicate abnormal vascular morphology alterations in the monkey hippocampus after 10 weeks of hTau overexpression. Many vascular abnormalities such as string vessels (open arrows), vessel occlusion (open arrowheads), as well as tortuosity, stenosis and bulging of the vascular walls (closed arrowheads) emerged after hTau overexpression. Scale bar, 50 μm. **(C)** Representative images of PECAM-1 immunostaining suggest vascular degeneration in the monkey hippocampus after hTau overexpression. There are significantly more disintegrated blood vessels in the monkey hippocampus after 6 weeks of hTau overexpression. Insets show higher magnification images of the blood vessels in the hippocampus. Scale bar, 1 mm (left) and 500 μm (inset, right). **(D)** Representative images of PECAM-1 immunostaining suggest vascular damages in the monkey hippocampus after hTau overexpression. The progressive vascular degeneration is mainly characterized by blood vessel ruptures (arrows) and disintegration (arrowheads). Scale bar, 20 μm. **(E)** Representative images of AQP4 and GFAP staining show relatively diffused AQP4 expression and substantial astrocytic endfeet retraction from blood vessels in the monkey hippocampus after hTau overexpression. Normally, astrocytic endfeet wrap blood vessels (open arrowheads) with AQP4 water channels on astrocytic endfeet forming ring-like structures (closed arrowheads). After tau overexpression, AQP4 signals become relatively diffused (closed arrows), and AQP4^+^ endfeet detach from the blood vessels (open arrows). Scale bar, 20 μm. **(F)** Quantification of the percentage of blood vessels with basement membrane thickening or distinct morphological changes in the monkey hippocampus. The percentages of basement membrane thickening or morphologically abnormal blood vessels in the monkey hippocampus after hTau overexpression are significantly higher than the percentages in the control monkey hippocampus. **P < 0.01. One-way ANOVA with Tukey’s post-hoc test. N = 3. **(G)** Quantification of the percentage of blood vessels with significant structural damages and disintegration in the monkey hippocampus. The percentages of damaged or degenerated vessels in the monkey hippocampus after hTau overexpression are significantly higher than the percentages in the control monkey hippocampus. *P < 0.05; ***P < 0.001. One-way ANOVA with Tukey’s post-hoc test. N = 3.

### 3.8. Learning and memory deficits and fine motor skill impairments after human tau overexpression

Cognitive decline, especially memory impairment, is a typical symptom of AD, and reduced hand dexterity has also been observed in individuals with AD (Buchman and Bennett 2011; de Paula, et al. 2016). Therefore, we utilized the Wisconsin General Test Apparatus (WGTA) to perform “delayed response” (DR) task (Supplementary video 1) and “delayed matching-to-sample/delayed nonmatching-to-s ample” (DMTS/DNMTS) task (Supplementary video 2-3) to test the spatial working memory and the associative learning ability of the rhesus monkeys before and after hTau overexpression (Rodriguez and Paule 2009; Tsutsui, et al. 2016). In addition, we also employed the rotating Brinkman board test (Supplementary video 4) to assess the finger fine motor skill of the same group of monkeys before and after hTau overexpression (Schmidlin, et al. 2011).

Six weeks after AAV injection, all the monkeys displayed an average decline of 50~60% in their spatial working memory capacity (how long the spatial information can be kept active in working memory, Fig. 8A-B). Furthermore, all the monkeys showed a profound impairment in the associative learning ability (how fast the monkeys can learn about the relationship between two separate stimuli) in two different DMTS/DNMTS-based associative learning tests (using Katakana and LEGO^®^ blocks as visual cues) after 8 weeks of hTau overexpression. On average, the monkeys would need roughly 50~70% more training time to acquire the skill to reliably retrieve (success rate ≥ 90%) the food rewards associated with certain visual cues (Fig. 8C-D and Supplementary Fig. 7A). Lastly, 10 weeks after viral injection, the food retrieval speed of most monkeys became slightly slower (~10%), whereas the food retrieval accuracy was relatively constant over the course of hTau overexpression, suggesting slightly deteriorated dexterity at a later stage of the pathology progression in the NHP disease models (Fig. 8E-F and Supplementary Fig. 7B).

**Figure 8.**
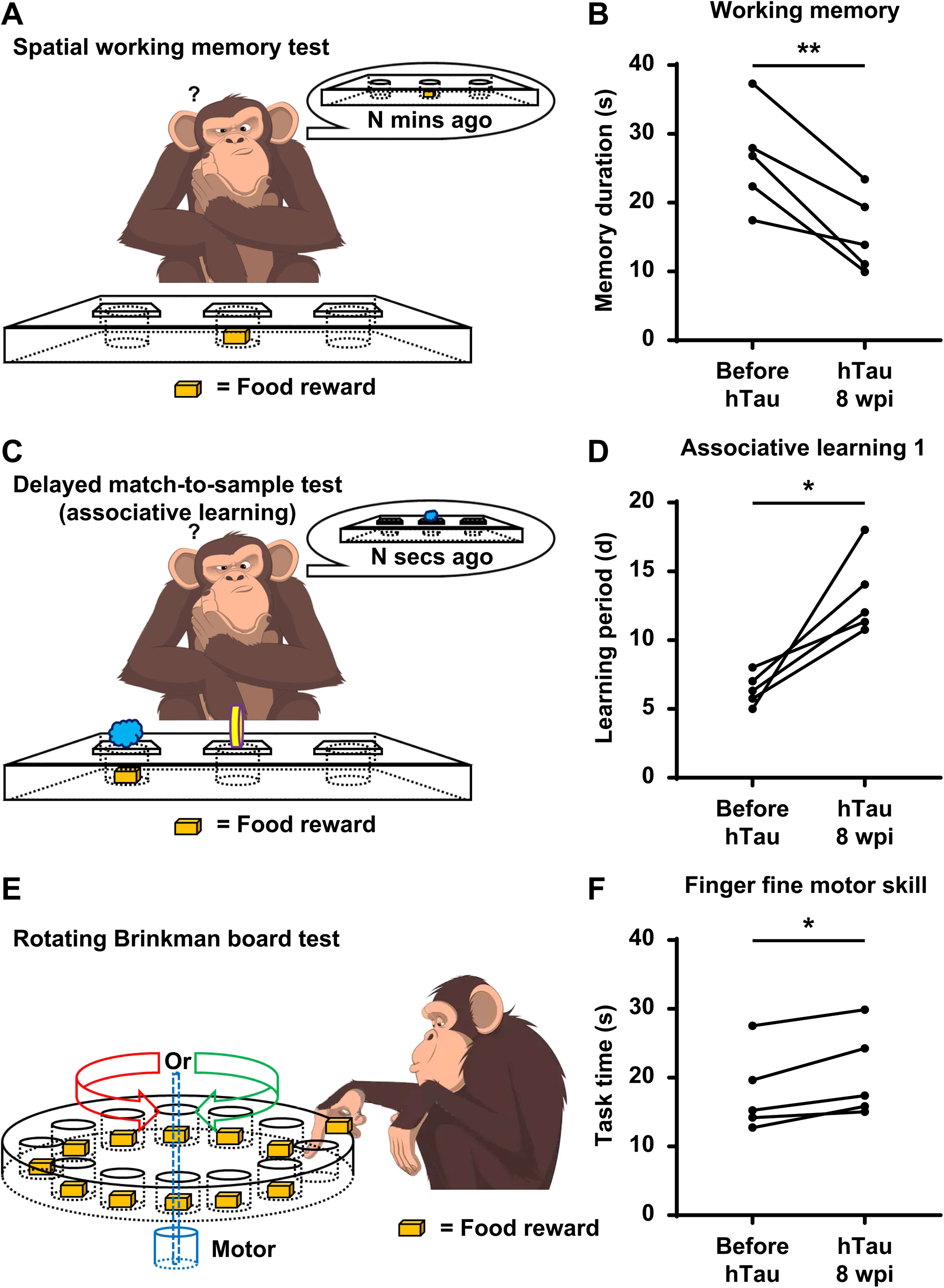
Deteriorated spatial working memory, impaired associative learning ability and compromised fine motor skill in monkeys with hTau overexpression. **(A)** A schematic drawing illustrates the experimental design of the WGTA-based monkey spatial working memory test. **(B)** Quantification of the memory duration before and after hTau overexpression. The working memory capacity of the monkeys declines significantly after hTau overexpression. Each dot represents an individual animal. **P < 0.01. Student’s t-test. N = 5. **(C)** A schematic drawing illustrates the experimental design of the WGTA-based monkey delayed match-to-sample test. **(D)** Quantification of the associative learning period before and after hTau overexpression. The learning ability is noticeably impaired in the monkeys after hTau overexpression. Each dot represents an individual animal. *P < 0.05. Student’s t-test. N = 5. **(E)** A schematic drawing illustrates the experimental design of the Brinkman board-based finger fine motor skill test. **(F)** Quantification of the food retrieval time before and after hTau overexpression. The monkeys’ food retrieval speed becomes slightly slower after hTau overexpression. Each dot represents an individual animal. *P < 0.05. Student’s t-test. N = 5.

Taken together, our AD-like NHP model can be generated via a single AAV injection within a couple of months, yet they recapitulate many defining pathological features of AD, including neuronal loss, hippocampal atrophy, inflammatory responses, Aβ clearance deficit, NFTs formation, vascular abnormalities and multimodal cognitive dysfunctions.

## 4. Discussion and conclusions

In this study, we establish an NHP model with a wide variety of Alzheimer’s-like pathologies ranging from the AD-related protein aggregation at the subcellular level to the AD-like memory deficits at the system level. The fact that it can be generated through a single stereotaxic injection of AAVs within 2 months may also help our model to achieve broader adoption within the biomedical research community.

NHP models of AD are important to bridging the translational gap between the findings from preclinical work in AD rodent models and effective therapies for AD patients. Through the collective efforts of the international community of Alzheimer’s researchers, some spontaneous, non-genetically or genetically induced NHP models of AD with great potential have been developed over the last two decades (Arnsten, et al. 2019; Beckman, et al. 2021; Forny-Germano, et al. 2014; Geula, et al. 2002; Geula, et al. 1998; Giannakopoulos, et al. 1997; Melamed, et al. 2017; Park, et al. 2015; Sani, et al. 2003; Sasaguri, et al. 2020; Seita, et al. 2020; Zhai, et al. 2018). However, most of these models take a very long time period and a substantial amount of cost to generate, rendering it difficult to conduct large-scale evaluations of therapeutic interventions over a practical time frame. In contrast, our NHP model with Alzheimer’s-like pathology can be conveniently produced via a single injection of AAVs into the brains of mid-aged rhesus monkeys within 2 months, and thus can be broadly adopted by the vast majority of ordinary researchers around the world to identify and analyze potential treatment strategies for AD within a reasonable timeline.

Until the relatively recent detection of p-tau tangles in their entorhinal cortex (Datta, et al. 2021; Paspalas, et al. 2018), rhesus macaques were not considered to be able to develop pathological tauopathy due to a lack of reporting of p-tau accumulation in their brains (Peters, et al. 1996; Zhang, et al. 2019). Since very few rhesus monkeys live long enough to reach 25+ years of age to show spontaneous tauopathy (Datta, et al. 2021; Paspalas, et al. 2018), induction of p-tau may become highly relevant in the future development of macaque AD models. In fact, leading researchers in the field have already been working on tauopathy induction models in the rhesus monkeys (Beckman, et al. 2021). Besides that, rhesus monkeys have rarely shown distinctive neuronal loss in their brains (Stonebarger, et al. 2021) or rapid deterioration of cognitive functions like AD patients (Hara, et al. 2012). Taking all the above into consideration, this study reveals an unexpected yet striking resemblance between our Alzheimer’s-like NHP model and Alzheimer’s patients. More specifically, our model not only exhibits widespread tauopathy and subsequent neuronal loss throughout the hippocampus, but also displays multimodal cognitive dysfunctions, neuroinflammation and microvascular alterations that characterize Alzheimer’s disease.

Excessive accumulations of Aβ and tau are widely recognized as the defining hallmarks of Alzheimer’s disease. Furthermore, mounting experimental and clinical evidences seem to point to a co-pathogenic interaction between Aβ and tau, which jointly accelerate the onset and progression of AD (Busche and Hyman 2020; Murphy and LeVine 2010). In line with this Aβ-tau synergetic theory, our Alzheimer’s-like NHP model induced by hTau overexpression in monkey hippocampi did display scattered intracellular Aβ accumulation, occasional parenchymal Aβ plaque-like deposits and significant Aβ clearance deficit revealed by immunostaining and CSF Aβ measurements. However, the distribution of Aβ aggregation in the brains of this NHP model is relatively sparse, which is inconsistent with the high density of senile plaques in many Alzheimer’s patients. Such discrepancy may be due to the long period of time that the abundant Aβ plaques need to build up through pTau-driven Aβ production or tau-related Aβ1-42 clearance (Arnsten, et al. 2021; Lonskaya, et al. 2014; Small, et al. 2017). We are planning to perform long-term measurement of Aβ deposition in the brains of our NHP model of AD to test this hypothesis in future studies.

Pathological studies of the brains from AD patients demonstrated that Alzheimer’s starts in the entorhinal cortex and hippocampus (Du, et al. 2001; Khan, et al. 2014), which is the exact reason why we chose to overexpress hTau in the hippocampi of the rhesus monkeys. However, Alzheimer’s disease will eventually lead to wide-spread damage across the whole brain, so an ideal NHP model of AD should display the major pathologic features not only in the hippocampus but also throughout the whole brain. Interestingly, we did detect robust overexpression of tau across the cerebral cortex of the monkeys 50 weeks after viral injection (Supplementary Fig. 2), which can probably be explained by the “prion-like” propagation of tau aggregation (Colin, et al. 2020; Mudher, et al. 2017). Since mounting evidence suggests that the propagation of tau would cause dysfunction and degeneration of the neuronal networks (Kaufman, et al. 2016; Stancu, et al. 2015), we will subsequently examine whether there is significant neuronal loss, neuroinflammation or microvascular abnormalities in all these cortical regions with tau pathology.

Hopefully, our AD-like NHP model with neuronal loss, hippocampal atrophy, neuroinflammation, Aβ clearance deficit, neurofibrillary tangle formation, blood vessel damage and cognitive decline will serve as an effective tool in further unveiling pathogenic mechanisms and developing disease-modifying treatments for AD in the foreseeable future.

## Acknowledgements

This work is supported by the National Natural Science Foundation of China (Grant No. 31970906), the Guangdong Provincial Natural Science Foundation (Grant No. 2020A1515010083), the Guangdong Grant ‘Key Technologies for Treatment of Brain Disorders’ (Grant No. 2018B030332001), and the Guangzhou Grant ‘Direct Astrocyte-to-Neuron Conversion to Treat Brain Disorders in NHPs’ (Grant No. 202206060002).

