## Supplementary Figure & Video Legends for "A non-human primate model with Alzheimer’s disease-like pathology induced by hippocampal overexpression of human tau"

Or

Gong Chen,.

1 These authors contributed equally to this work.

**Supplementary figure 1.** Representative images of tau immunostaining demonstrate the AAV-induced overexpression of tau in neurons within the monkey hippocampus 10 weeks after viral injection. Compared to that of endogenous tau (top), the expression level of tau in the hTau overexpressed monkey brains (bottom) is significantly higher. Scale bar, 1 mm.

**Supplementary figure 2.** Representative images of tau immunostaining show the overexpression of tau throughout the monkey brain 50 weeks after viral injection, indicates tau pathology spreading from the AAV injection sites to the whole brain possibly through prion like propagation. Insets show higher magnification of the hippocampal (a1, b1) and cortical (a2, b2, a3, b3, a4, b4) regions. Scale bar, 1 mm (half brain, left) and 50 μm (inset, bottom right).

**Supplementary figure 3.** Representative images of NeuN and tau immunostaining indicate neuronal degeneration and loss within the monkey hippocampus 6 weeks after viral injection. The number of NeuN^+^ cells decreases significantly wherever the hTau expression level is high. Insets show higher magnification of the CA3 (a1, b1) and CA1 (a2, b2) regions. Scale bar, 1 mm (top) and 20 μm (inset, bottom).

**Supplementary figure 4.** Representative images of GFAP immunostaining show astrocytic activation within the monkey hippocampus. The number of GFAP^+^ cells increases significantly in the monkey hippocampus after 6 weeks of hTau overexpression. Meanwhile, many GFAP^+^ cells in the hippocampus undergo typical morphological changes seen in reactive astrocytes after hTau overexpression. Insets show higher magnification of the CA3 (a1, b1) and CA1 (a2, b2) regions. Scale bar, 1 mm (top) and 20 μm (inset, bottom).

**Supplementary figure 5.** Representative images of Iba1 immunostaining suggest microglial activation and neuroinflammatory response in the monkey hippocampus. The number of Iba1^+^ cells increases significantly in the monkey hippocampus after 6 weeks of hTau overexpression. Note that many Iba1^+^ cells in the hippocampus undergo typical morphological changes seen in reactive microglia after hTau overexpression. Insets show higher magnification of the CA3 (a1, b1) and CA1 (a2, b2) regions. Scale bar, 1 mm (top) and 20 μm (inset, bottom).

**Supplementary figure 6.** Representative images of CD68 immunostaining show elevated CD68 signals across the hippocampus, further confirm robust immune activation and neuroinflammation throughout the hippocampus after 6 weeks of hTau overexpression. Insets show higher magnification of the CA3 region. Scale bar, 1 mm (top) and 20 μm (inset, bottom).

**Supplementary figure 7. Results of some other NHP behavioral tests. (A)** Quantification of the associative learning period before and after hTau overexpression. The learning ability is noticeably impaired after 8 weeks of hTau overexpression. Each dot represents an individual animal. *P < 0.05. Student's t-test. N = 5. **(B)** Quantification of the success rate of food retrieval before and after hTau overexpression. The success rate of food retrieval remains unchanged after 8 weeks of hTau overexpression. Each dot represents an individual animal. “n.s.” means “not (statistically) significant” (P>0.05). Student's t-test. N = 5.

**Supplementary video 1. WGTA-based monkey spatial working memory test.** Delayed response (DR) task was conducted using the Wisconsin General Test Apparatus (WGTA), in which a monkey sat in its home cage in front of a tray that contains 3 food wells covered by identical swing-away lids. Initially, the experimenter baited one of the wells with food in front of the monkey, covered it, and then lowered an opaque screen to block the food tray from the monkey’s view. After a certain delay period the screen was removed, which allowed the monkey to retrieve the food from the baited food well which had to be recalled from working memory. Food reinforcers were randomly distributed among 3 wells over 30 trials per day (≥ 26 correct choices for 3 consecutive days to pass), and the length of the delays were gradually increased according to a stepwise procedure as the monkeys demonstrate mastery of the task.

**Supplementary video 2. WGTA-based monkey delayed matching-to-sample test.** Delayed matching-to-sample (DMTS) task was also performed with a WGTA. The experimenter first presented a sample visual stimulus (one of katakana characters or LEGO^®^ blocks) to the monkey. Following a randomly interval (up to 5 seconds), the same sample visual stimulus and another visual distractor were both presented to the monkey, and the monkey needed to choose the visual stimulus that matched the sample stimulus to get the food reward. Each incorrect choice was followed by a 10 seconds timeout. The monkeys took 30 DMTS trials per day (≥ 26 correct choices for 3 consecutive days to pass), and both the sample visual stimuli and the visual distractors were randomly selected each day from either 46 basic katakana alphabet letters or a large pool of LEGO^®^ blocks (> 200 combinations).

**Supplementary video 3. WGTA-based monkey delayed nonmatching-to-sample test.** Delayed nonmatching-to-sample (DMTS) task was also performed with a WGTA. The experimenter first presented a sample visual stimulus (one of katakana characters or LEGO^®^ blocks) to the monkey. Following a randomly interval (up to 5 seconds), the same sample visual stimulus and another visual distractor were both presented to the monkey, and the monkey needed to choose the visual stimulus that is different from the sample stimulus to get the food reward. Each incorrect choice was followed by a 10 seconds timeout. The monkeys took 30 DMTS trials per day (≥ 26 correct choices for 3 consecutive days to pass), and both the visual stimuli were randomly selected each day from either 46 basic katakana alphabet letters or a large pool of LEGO^®^ blocks (> 200 combinations).

**Supplementary video 4. Brinkman board-based monkey finger fine motor skill test.** Rotating Brinkman board task is a manual dexterity task for monkeys, in which 14 food pellets were placed in 14 narrow slots (60 mm length × 6 mm width × 6 mm depth) on a rotating brinkman board (240 mm in diameter), and the monkeys were trained to retrieve all the food pellets from the slots using one of their hands. Monkeys were trained to master this task for 3 consecutive days before they were tested each day for retrieval accuracy and speed for 5 consecutive days. During both the training and testing sessions, the monkeys performed 4 sessions each day, among which the board was rotating clockwise in 2 sessions and counterclockwise in the other 2 sessions, and the monkeys were allowed to only use their left hands in 2 sessions and right hands in the other 2 sessions. All experimental procedures were recorded with FHD video cameras.
