## Supplementary figures and images for "A non-human primate model with Alzheimer’s disease-like pathology induced by hippocampal overexpression of human tau"

### Supplementary Figure_1

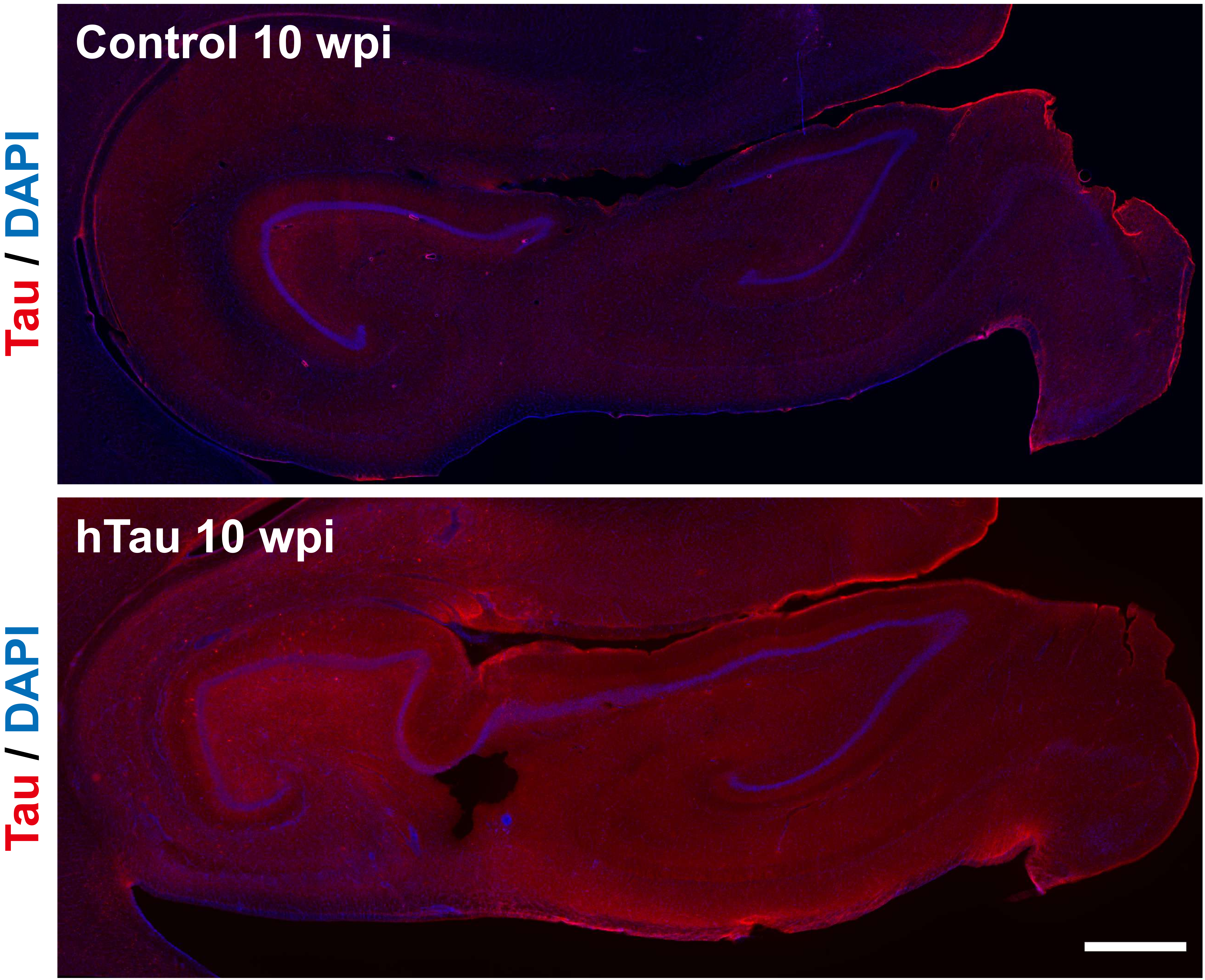

### Supplementary Figure_2

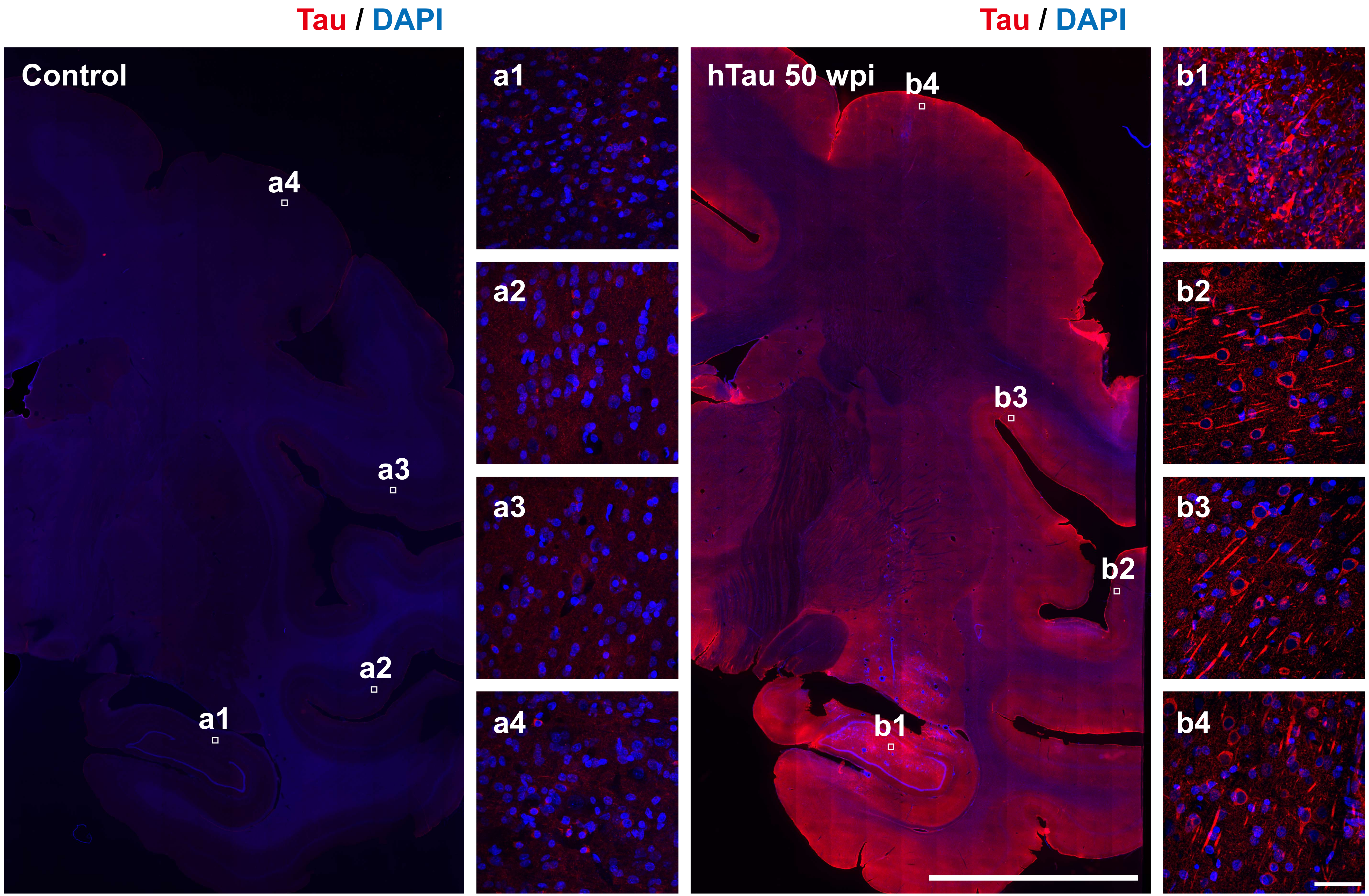

### Supplementary Figure_3

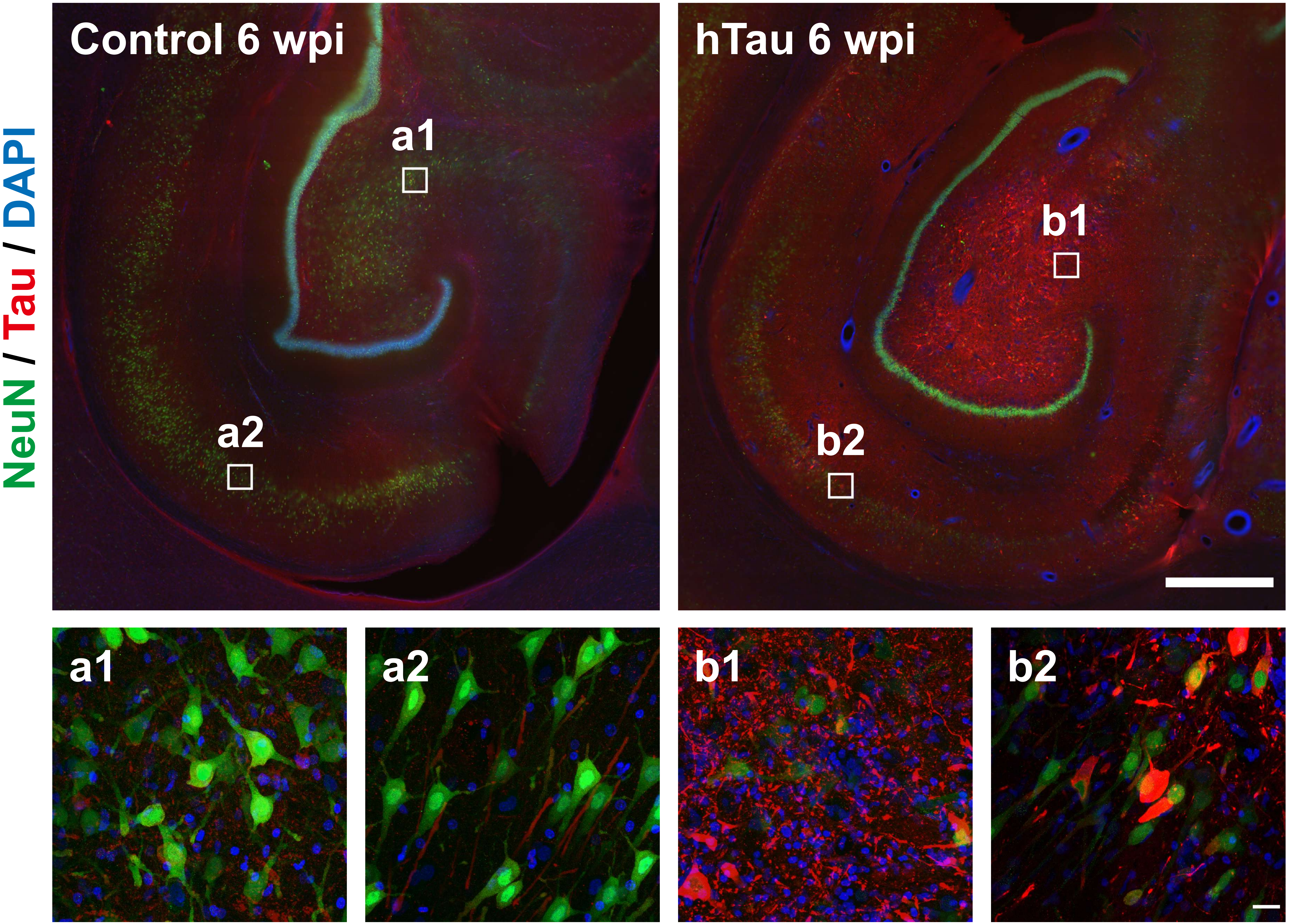

### Supplementary Figure_4

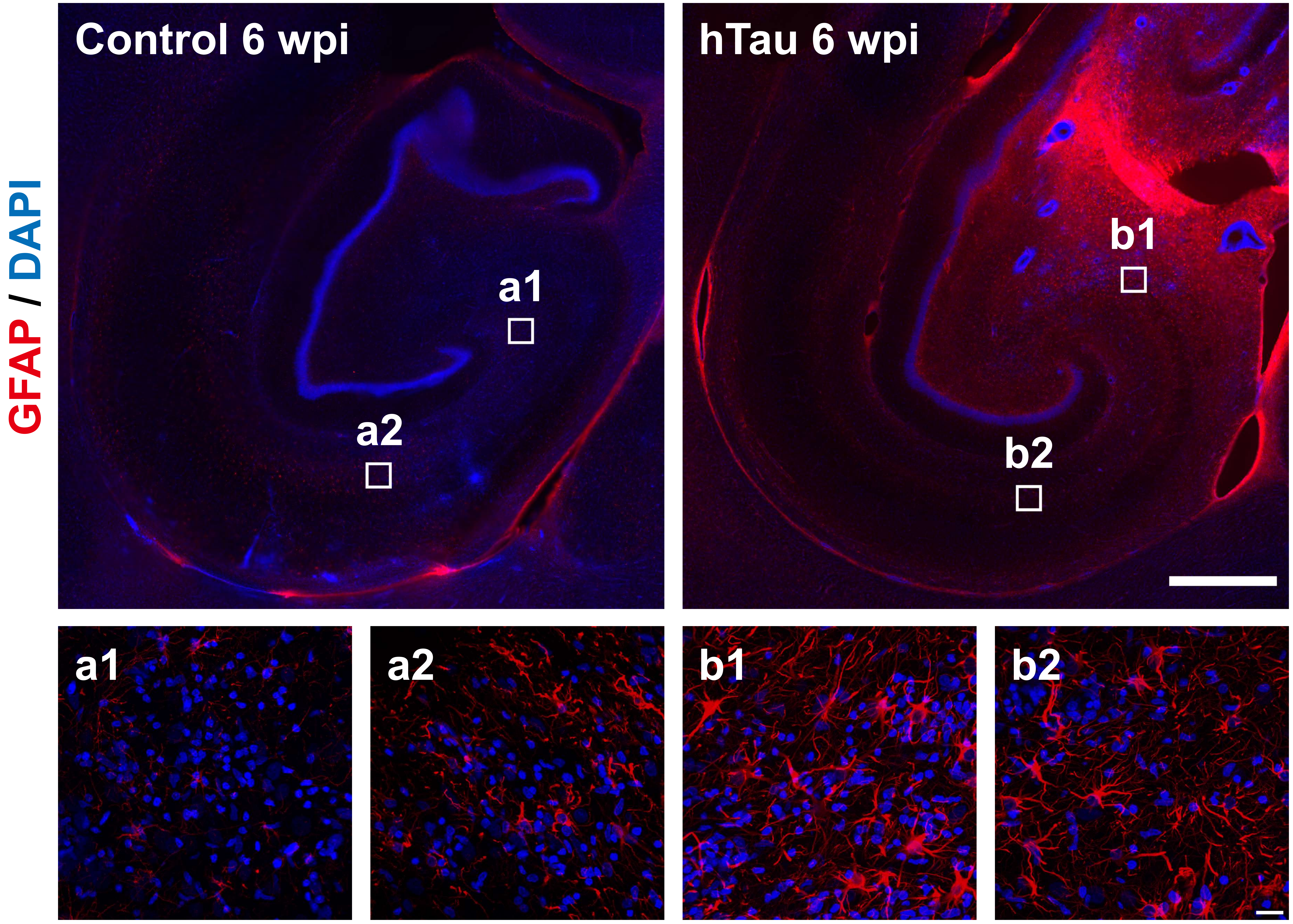

### Supplementary Figure_5

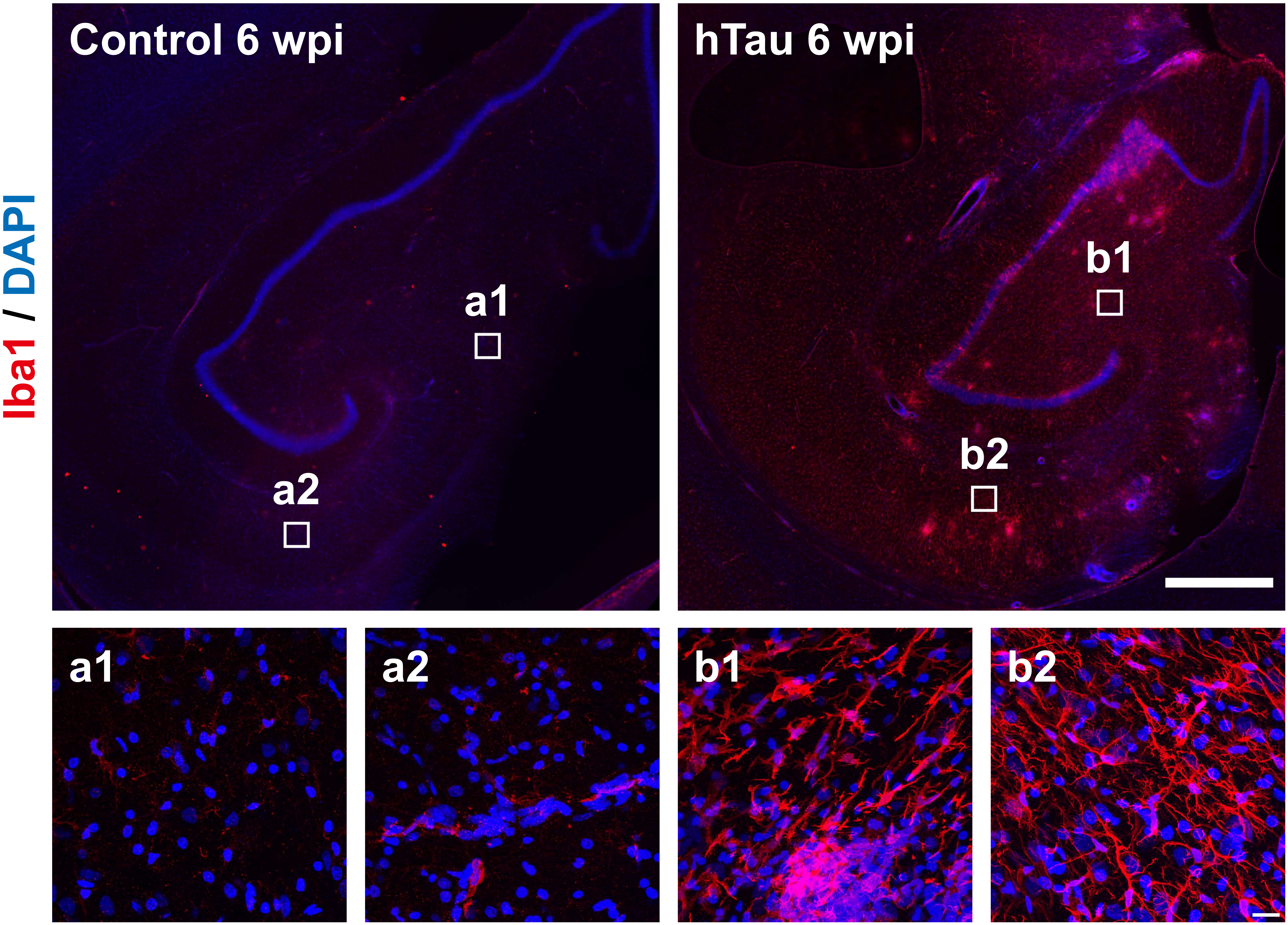

### Supplementary Figure_6

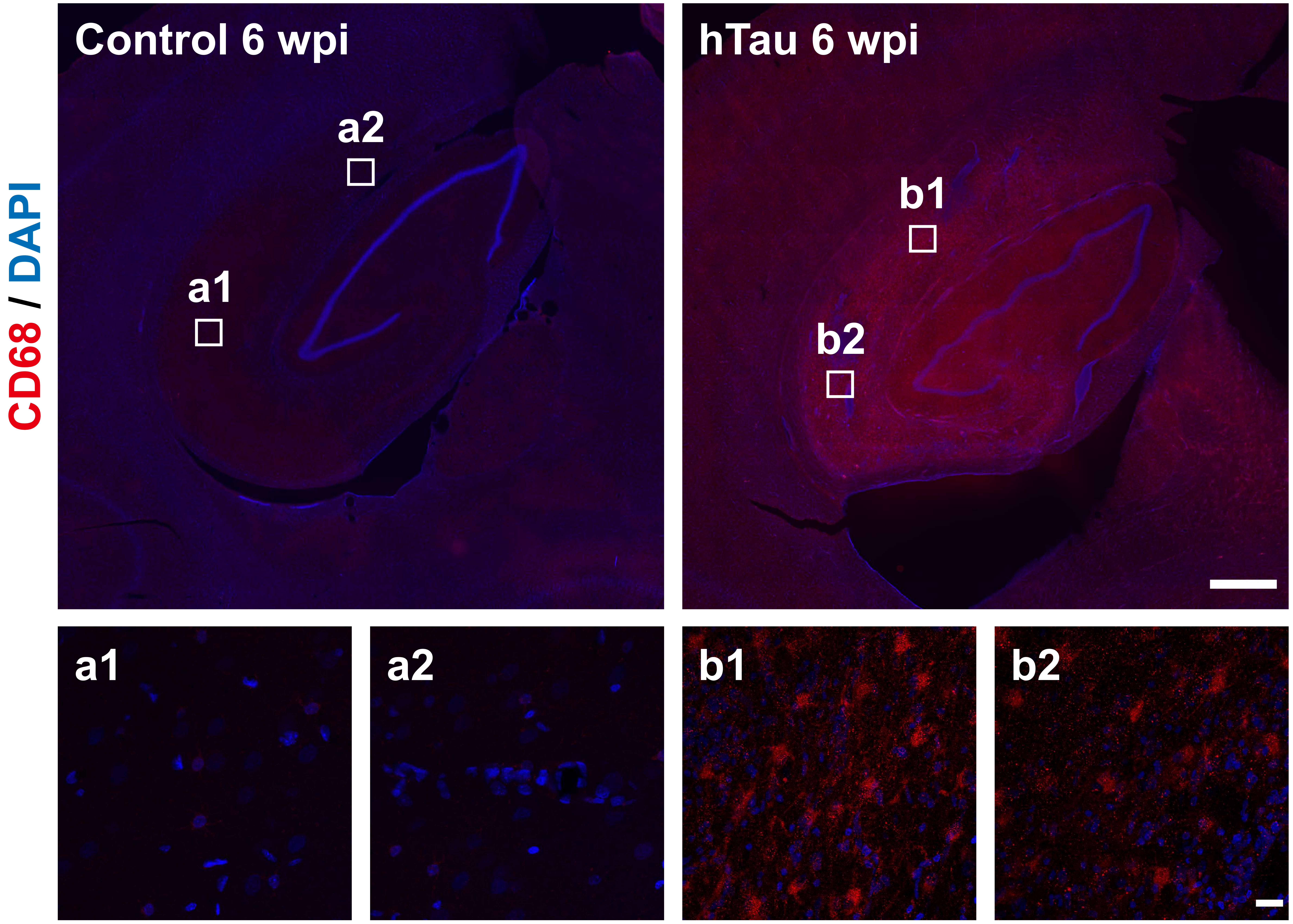

### Supplementary Figure_7

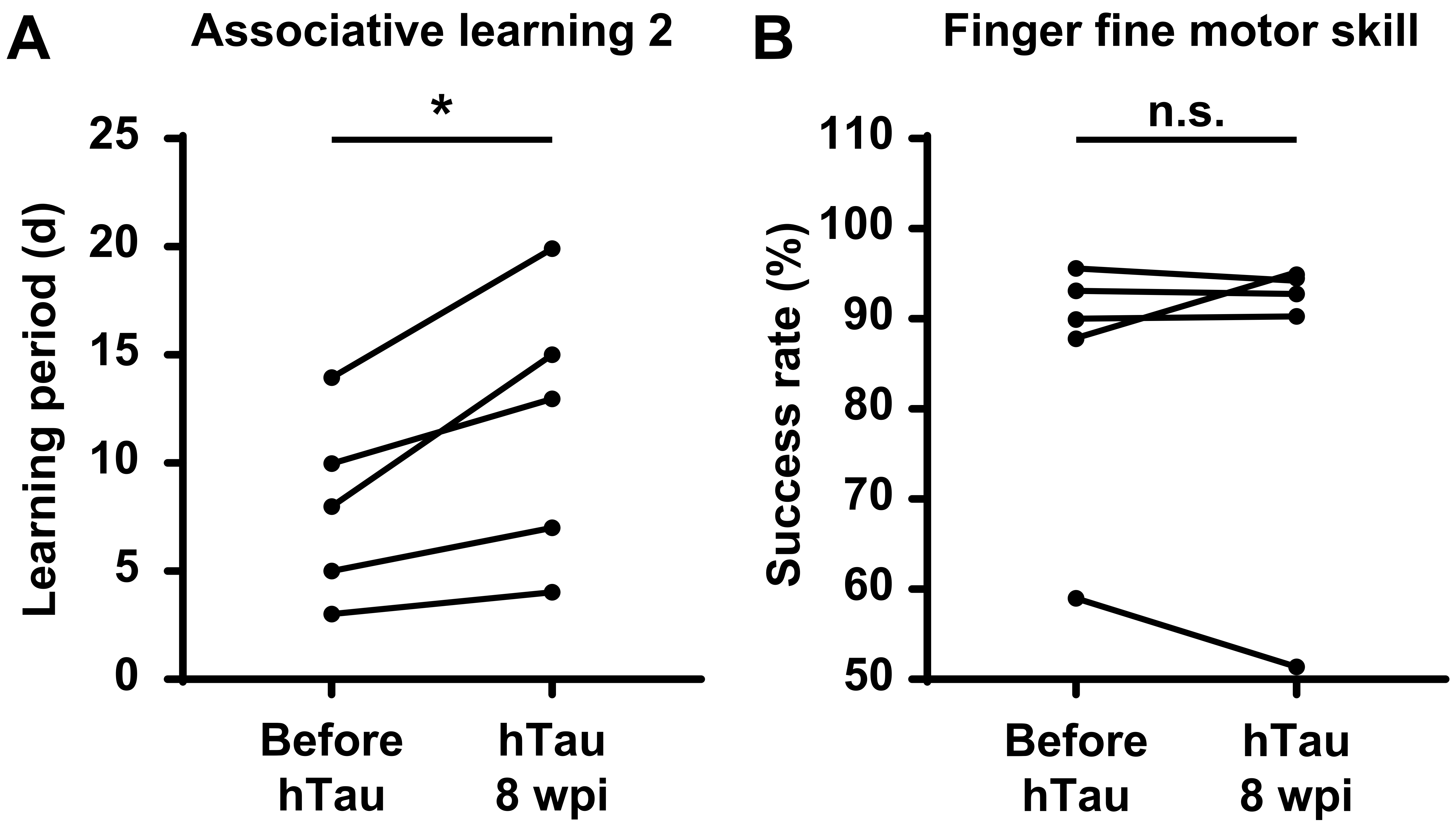
